# Visual pathways in the brain of the jumping spider *Marpissa muscosa*

**DOI:** 10.1101/640706

**Authors:** Philip O.M. Steinhoff, Gabriele Uhl, Steffen Harzsch, Andy Sombke

## Abstract

Some animals have evolved task differentiation among their eyes. A particular example is spiders, where most species have eight eyes, of which two (the principal eyes) are used for object discrimination, whereas the other three pairs (secondary eyes) detect movement. In the spider species *Cupiennius salei* these two eye types correspond to two visual pathways in the brain. Each eye is associated with its own first and second order visual neuropil. The second order neuropils of the principal eyes are connected to the arcuate body, whereas the second order neuropils of the secondary eyes are linked to the mushroom body. However, eye size and visual fields are considerably different in jumping spiders. We explored the principal- and secondary eye visual pathways of the jumping spider *Marpissa muscosa*. We found that the connectivity of the principal eye pathway is the same as in *C. salei*, while there are differences in the secondary eye pathways. In *M. muscosa,* all secondary eyes are connected to their own first order visual neuropils. The first order visual neuropils of the anterior lateral and posterior lateral eyes are further connected with two second order visual neuropils, whereas the posterior median eyes lack second order visual neuropils and their axons project only to the arcuate body. This suggests that the posterior median eyes probably do not serve movement detection in *M. muscosa.* Furthermore, the second order visual neuropil (L2) in *Marpissa muscosa* potentially integrates information from the secondary eyes and might thus enable faster movement decisions.

## 1 Introduction

In some animal species with multiple eyes, different eyes serve different specific tasks, such as compound eyes and ocelli in insects or the rhopalia in box jellyfishes (Garm et al., 2007; O’Connor et al., 2009; Paulus, 1979). Spiders vary strongly in their visual abilities, as well as in number, arrangement and size of their eyes (reviewed in Morehouse et al. 2017). Most species possess four pairs of eyes, of which one pair differs from the three other pairs in its anatomy and developmental origin (Land, 1985a; Morehouse et al., 2017). The principal eyes (or anterior median eyes) possess a movable retina and their spectral sensitivities allow for color vision (Barth et al. 1993; Land, 1985b; Schmid, 1998; Yamashita & Tateda, 1976, 1978). The other three pairs of eyes are the so-called secondary eyes (posterior median, anterior lateral and posterior lateral eyes). Their morphology and anatomy differs considerably among spider families (Homann, 1952, 1950), but they generally do not have a movable retina. In several groups of cursorial hunting spiders, the secondary eyes are used for movement detection (Land, 1985a; Morehouse et al., 2017; Schmid, 1998). Most secondary eyes possess a light-reflecting tapetum (but absent in e.g. jumping spiders), which led to the assumption that they play a major role for night or dim light vision (Foelix, 1996).

Jumping spiders (Salticidae) are cursorial hunters with excellent vision. Their eyes have been studied in much greater detail compared to any other spider group (e.g. Harland et al., 2012; Land, 1985a; Morehouse et al., 2017). The principal eyes of jumping spiders are large and face forward; their field of view is small, but their spatial acuity is one of the highest among invertebrate eyes (Harland et al., 2012; Land, 1985b, 1985a). The secondary eyes of salticids possess only one photoreceptor type and thus cannot discriminate colors (Yamashita and Tateda, 1976). Among the secondary eyes, the anterior lateral eyes (ALE) and posterior lateral eyes (PLE) are large, able to detect motion, and elicit an orienting response of the principal eyes towards the moving object (Duelli, 1978; Jakob et al., 2018; Spano et al., 2012; Zurek et al., 2010; Zurek and Nelson, 2012a, 2012b). The posterior median eyes (PME) of most salticids (except for the non-salticoids) are very small and positioned between ALE and PLE (Land, 1985b; Maddison and Hedin, 2003). Since the fields of view of ALE and PLE overlap, the PME do not have an exclusive field of view (Land, 1985b).

While the eyes of jumping spiders have been studied in detail, less is known about the brain structures that process visual information (Duelli, 1980; but see Hanström, 1921; Hill, 1975; Nagata et al., 2019; Steinhoff et al., 2017; Strausfeld, 2012). Our knowledge on the visual processing pathways in the brain of spiders is mostly based on investigations of the spider model species *Cupiennius salei* (Keyserling, 1877) (Babu and Barth, 1984; Becherer and Schmid, 1999; Schmid and Duncker, 1993; Strausfeld et al., 1993; Strausfeld and Barth, 1993). *C. salei* possesses two separate visual pathways, one for the principal eyes and one for all secondary eyes (Strausfeld et al., 1993; Strausfeld and Barth, 1993). In *C. salei*, every eye is connected to its own first and second order visual neuropil (Barth, 2002). While the second order visual neuropils of the principal eyes are connected to the arcuate body (Strausfeld et al., 1993), the second order visual neuropils of the secondary eyes are associated with the mushroom bodies (Strausfeld and Barth, 1993). Additionally, the first order visual neuropils of the secondary eyes are directly connected to the arcuate body (Babu and Barth, 1984). In *C. salei,* similar to salticids, the principal eyes are used for object recognition, while the secondary eyes serve the purpose of movement detection (Fenk, Hoinkes, & Schmid, 2010; Schmid, 1998). The secondary eyes of *C. salei* have a high spectral sensitivity, similar to their principal eyes (Barth et al., 1993) and similar to jumping spiders are capable of acute motion detection (Fenk & Schmid, 2010; Zurek & Nelson, 2012b). However, they are unable to detect color in moving objects (Orlando and Schmid, 2011) and the relative size of different eyes and thus fields of view in *C. salei* differ considerably from jumping spiders (Land, 1985b; Land and Barth, 1992). Furthermore, Steinhoff et al. (2017) showed that the structure and arrangement of the secondary eye visual neuropils in the jumping spider *Marpissa muscosa* (Clerck, 1757) differ from those in *C. salei.* We thus hypothesize, that the secondary eye visual pathways of jumping spiders also differ from that in the model species *C. salei.* Here, we describe the principal and secondary eye visual pathways in the brain of the jumping spider *M. muscosa*. We use paraffin histology, immunohistochemistry, and microCT analysis to visualize tracts in the visual system. We compare our findings in *M. muscosa* with results of earlier studies and new immunohistochemical data on the principal and secondary eye pathways in the brain of *C. salei*. We discuss the similarities and differences in the principal and secondary eye pathways of these two spider species in the light of different functional roles of their eyes.

## 2 Material and Methods

### 2.1 Animals

Adult female *Marpissa muscosa* were collected in and near Greifswald (Germany). Spiders were fed on *Drosophila* sp. weekly and kept individually in plastic boxes of 145 × 110 × 68 mm size that were enriched with paper tissue.

### 2.2 Paraffin-histology

The prosomata of two females of *M. muscosa* were fixed in Gregory’s artificially aged Bouin solution (Gregory, 1980). After four days in fixative, the prosomata were dehydrated through a graded ethanol series (80%, 90%, and 3× 96% ethanol for 20 minutes each) and were then transferred for two hours into a 1:1 solution of 96% ethanol:tetrahydrofuran (Carl Roth, CP82.1) at room temperature. Samples were kept for 24 hours in pure tetrahydrofuran followed by 24 hours in a solution of 1:1 tetrahydrofuran and paraffin (Carl Roth, 6643.1) at 60°C. Afterwards, samples were transferred to 100% paraffin at 60°C for 48 hours and then embedded in fresh paraffin. Sagittal and transversal sections (5 μm) were cut with a motorized rotary microtome (Microm HM 360). Sections were stained with Azan according to Geidies (Schulze and Graupner, 1960), and mounted in Roti-Histokitt II (Carl Roth, T160.1).

### 2.3 MicroCT analysis

Prosomata of two individuals of *M. muscosa* and one individual of *C. salei* were fixed in Dubosq–Brazil solution (1 g saturated alcoholic picric acid, 150 ml 80% ethanol, 15 ml pure acetic acid, and 60 ml 37% formaldehyde). Prior to fixation, legs and opisthosomata were cut off and cuticle and musculature were removed from the dorsocaudal part of the prosomata to allow faster penetration of the fixative. After four days in fixative, samples were washed six times for 20 minutes in 0.1 M phosphate buffered saline (PBS, pH 7.4) followed by dehydration in a graded ethanol series (80%, 90%, 96% and 3x 99,8% ethanol for 20 minutes each). Samples were then transferred to an 1% iodine solution (iodine, resublimated [Carl Roth, X864.1] in 99.8% ethanol) over 48 h to enhance tissue contrast (Sombke et al., 2015) and subsequently critical point dried with an automated dryer (Leica EM CPD300). The protocol applied was: slow CO_2_ admittance with a delay of 120 seconds, 30 exchange cycles, followed by a slow heating process and slow gas discharge. Dried prosomata were mounted using a conventional glue gun onto aluminum rods, so that the anterior median eyes were oriented upwards.

MicroCT scans were performed using a Zeiss XradiaXCT-200 (Sombke et al., 2015). Scans were performed with a macro-, a 4x- and a 20x objective lens unit and the following settings: 40 kV and 8 W or 30 kV and 6 W, 200 μA, exposure times between 2 and 4 s. Tomographic projections were reconstructed with the XMReconstructor software (Zeiss), resulting in image stacks (TIFF format). Scans were performed using Binning 2 for noise reduction and were reconstructed with full resolution (using Binning 1).

### 2.4 Immunohistochemistry

Four different combinations of markers were used to visualize neuropils and connecting neurites in the visual system (see Table 1 for a list of marker combinations used, and Table 2 for specifications of labeling reagents).

**Table 1.**
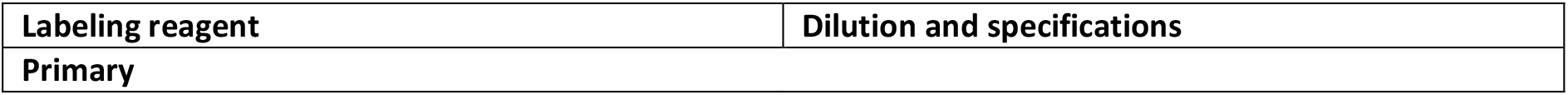
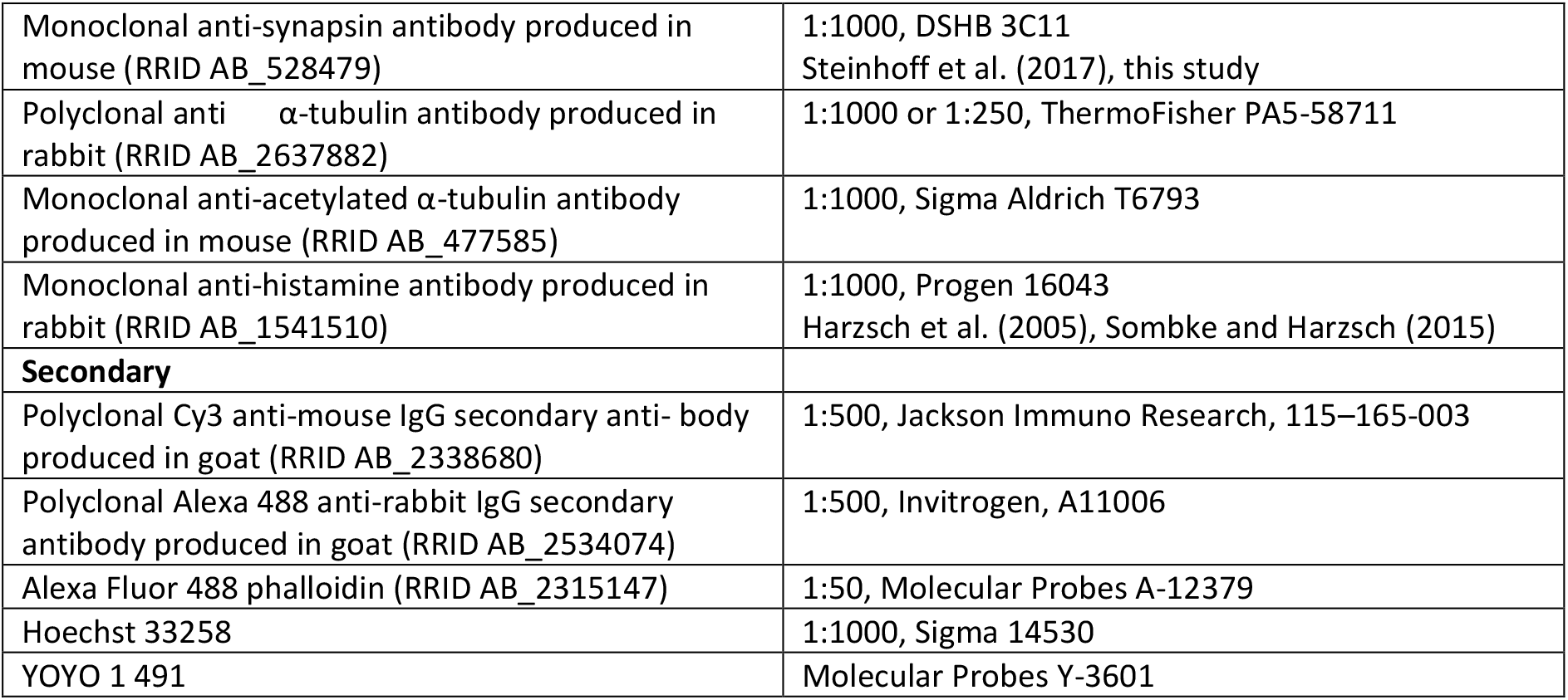
Primary and secondary antibodies

| Labeling reagent | Dilution and specifications |
| --- | --- |
| Primary |  |
| Monoclonal anti-synapsin antibody produced in mouse (RRID AB_528479) | 1:1000, DSHB 3C11<br>Steinhoff et al. (2017), this study |
| Polyclonal anti-α-tubulin antibody produced in rabbit (RRID AB_2637882) | 1:1000 or 1:250, ThermoFisher PA5-58711 |
| Monoclonal anti-acetylated α-tubulin antibody produced in mouse (RRID AB_477585) | 1:1000, Sigma Aldrich T6793 |
| Monoclonal anti-histamine antibody produced in rabbit (RRID AB_1541510) | 1:1000, Progen 16043<br>Harzsch et al. (2005), Sombke and Harzsch (2015) |
| **Secondary** |  |
| Polyclonal Cy3 anti-mouse IgG secondary anti-body produced in goat (RRID AB_2338680) | 1:500, Jackson Immuno Research, 115-165-003 |
| Polyclonal Alexa 488 anti-rabbit IgG secondary antibody produced in goat (RRID AB_2534074) | 1:500, Invitrogen, A11006 |
| Alexa Fluor 488 phalloidin (RRID AB_2315147) | 1:50, Molecular Probes A-12379 |
| Hoechst 33258 | 1:1000, Sigma 14530 |
| YOYO 1 491 | Molecular Probes Y-3601 |

**Table 2.** Methods and antisera combinations

| Methods | Antisera combinations/ staining | Coupled with | Specimens |
| --- | --- | --- | --- |
| Wholemount | anti-synapsin | Cy3 anti-mouse (goat) | 5 |
| Vibratome sections<br>(50 μm; 80 μm) | anti-synapsin (mouse)<br>anti-α-tubulin (rabbit) | Cy3 anti-mouse (goat)<br>Alexa Fluor 488 anti-rabbit<br>(goat) Nuclear labeling: Hoechst<br>33258 | 16 |
| | anti-acetylated-α-tubulin (mouse) | Nuclear labeling: YOYO | 6 |
|  | anti-histamine (rabbit) | Alexa Fluor 488 anti-rabbit<br>(goat) | 4 |
|  | anti-synapsin (mouse) | Cy3 anti-mouse (goat)<br>Phalloidin | 5 |
| Paraffin sections<br>(5 μm) | Azan-novum | / | 2 |
| microCT scans | 1% iodine (in 99% ethanol) | / | 4 |

After anaesthetization with CO_2_, brains of 31 females of *M. muscosa* (details Tab. 2) and four females of *C. salei* were dissected in PBS and fixed in 4% paraformaldehyde over night at room temperature. Specimens for histamine immunohistochemistry were prefixed overnight at 4°C in 4% N-(3-Dimethylaminopropyl)-N’-ethylcarbodiimide hydrochloride (EDAC, Sigma–Aldrich E6383). Brains were washed six times for 15 minutes each in PBS and subsequently immersed in Poly-L-Lysin (Sigma-Aldrich, P4707) for one minute. Samples were then embedded in 4% agarose (Sigma-Aldrich, A9414) and sectioned (50 or 80 μm) with a Microm HM 650 V vibratome. After permeabilization in PBS-TX (PBS, 0.3% Triton X-100 [Sigma-Aldrich, X100], 1 % bovine serum albumin [Sigma-Aldrich, A2153], 0.05% Na-acide; or alternatively PBS, 0.5% Triton, 1.5% DMSO) for 1 h at room temperature, incubation in primary antibodies took place over night at 4°C or room temperature. Sections were washed in several changes of PBS-TX after the incubation. Incubation in secondary antibodies or counterstains took place overnight at 4°C or room temperature. After washing in several changes of PBS, sections were mounted in Mowiol (Merck 475904).

The brains of five additional females of *M. muscosa* were processed as whole-mounts, using an adjusted protocol after Ott (2008) (see also Steinhoff et al., 2017). Brains were dissected in TRIS buffer (0.1 M, pH 7.4 [Carl Roth, 4855]), and fixed in zinc-formaldehyde at room temperature under soft agitation for several days. Afterwards, brains were washed in TRIS buffer three times for 15 minutes each and incubated in a 4:1 mixture of methanol and DMSO (Dimethylsulfoxide [Carl Roth, 4720]) for 2 h. After incubation in 100% methanol for 1 h, brains were rehydrated in a graded series of methanol (90%, 70%, 50%, 30%) for 10 minutes each and washed in TRIS buffer. This was followed by permeabilization in PBS-DMSO (PBS, 5% bovine serum albumin, 1% DMSO, 0.05% Na-acide) for 2 h at room temperature and incubation in primary antibody for 4 days at 4°C. Brains were then washed several times for 2 h in PBS-DMSO and incubated in secondary antibody for 4 days at 4°C, and transferred to a graded series of glycerol, with 1%, 2%, 4% (for two hours each) and 8%, 15%, 30%, 50%, 60%, 70%, 80% (for one hour each) glycerol in TRIS buffer and 1% DMSO. After five changes in pure ethanol for 30 minutes each, brains were mounted in fresh methyl salicylate (Sigma, 76631). In control experiments, in which we replaced the primary antibodies with PBS-TX, no neuronal labeling was detected.

### 2.5 Western Blot

In western blots of *Drosophila melanogaster* Meigen, 1830 head homogenates (Klagges et al., 1996), the monoclonal antibody mouse anti-*Drosophila* synapsin ‘‘Synorf 1‘‘ (provided by E. Buchner, Universität Würzburg, Germany; raised against a *Drosophila* Glutathione S-Transferase(GST)-synapsin fusion protein) recognizes at least 4 synapsin isoforms (ca. 70, 74, 80, and 143 kDa). In western blot analyses in Crustacea (Harzsch and Hansson, 2008; Sullivan et al., 2007) and Chilopoda (Sombke et al., 2011), isoforms in the same range (75-90 kDA) were recognized. Furthermore, the Synorf 1 antibody labels synaptic neuropil in taxa as diverse as Araneae (Fabian-Fine et al., 1999; Nagata et al., 2019; Steinhoff et al., 2017; Widmer et al., 2005), Chaetognatha (Harzsch and Müller, 2007) and Plathelminthes (Cebrià, 2008), suggesting that the epitope that this antiserum recognizes is highly conserved. To test this, we conducted a western blot analysis as well, in which we compared brain tissue of *D. melanogaster* and *M. muscosa*. The antibody provided similar results for both species, staining one strong band around 70 kDa, and in *M. muscosa* another weak band around 60 kDa. This result suggests that the epitope that Synorf 1 recognizes is conserved between *D. melanogaster* and *M. muscosa*.

### 2.6 Microscopy, image processing and nomenclature

Immunohistochemically labelled sections and whole-mounts were examined and photographed with a Leica SP5 II confocal microscope (cLSM). Paraffin sections were examined and photographed with a customized Visionary Digital BK Plus Lab System (duninc.com/bk-plus-lab-system.html). 3D-visualization of microCT image stacks was prepared in AMIRA 6.0.0 (ThermoFisher). An interactive 3D-visualization of the microCT reconstructions was created using ImageJ 1.51n and Adobe Acrobat Pro 9.0 (see supporting information; Figure S1). Images were processed in Corel PaintShop Pro using global contrast and brightness adjustment features as well as black and white inversion. Schematic drawings were generated using Corel Draw 20.1 and figure plates were assembled in Corel PaintShop Pro 21.0. A color code for all depicted structures, as used in Loesel et al. (2013) and Steinhoff et al. (2017), was used for 3D visualizations and the schematic drawings. The terminology used for brain structures follows Richter et al. (2010). For structures specific to spiders, we use terms according to Lehmann et al. (2016), Strausfeld et al. (1993) and Strausfeld and Barth (1993). Positional information is given with respect to the body axis. We are referring to results from immunohistochemical experiments as immunoreactivity, e.g. synapsin-immunoreactivity.

#### Abbreviations

AB: Arcuate body
ALE: Anterior lateral eyes
AL1: First order visual neuropils of ALE
AL2/PL2: Second order visual neuropils of ALE and PLE
AL1x: Neuropilar subunit of the AL1
AME: Anterior median eyes
AM1: First order visual neuropils of AME
AM2: Second order visual neuropils of AME
L2: Second order visual neuropils of anterior and posterior lateral eyes
MB: Mushroom bodies (neuropil with attributes of a mushroom body-like organization)
MBbr: Mushroom body bridge
MBn: Neurites between pedunculus and AL2/PL2
MBh: Mushroom body haft
MBp: Pedunculus of the mushroom body
MBs: Mushroom body shaft
PLE: Posterior lateral eyes
PL1: First order visual neuropils of the PLE
PL1x: Neuropilar subunit of the PL1
PME: Posterior median eyes
PM1: First order visual neuropils of the PME
PM2: Second order visual neuropils of the PME
SC1-3: Protocerebral soma cluster 1-3

## 3 Results

### 3.1 General organization of the protocerebrum

In *Marpissa muscosa,* the protocerebrum comprises bilaterally paired neuropils of the visual system and the mushroom body as well as the unpaired arcuate body and the protocerebral neuropil (Figs. 1, 2; for details see Steinhoff et al., 2017). All neuropils of the visual system are recognizable in paraffin sections (Fig. 3) and microCT analysis (Fig. 4; supporting information, Fig. S1), and are also characterized by a strong synapsin-immunoreactivity (Figs. 1; 5; 6; 7a, b). Tracts and individual neurites exhibit prominent tubulin-immunoreactivity (Figs. 1; 2; 5; 6; 7). Furthermore, the optic nerves (between retinae and first order visual neuropils) and the corresponding first order visual neuropils display histamine-immunoreactivity (Fig. 8), which is known to consistently label retinula cell axons and their terminals in the visual systems of arthropods (Battelle et al., 1991; Harzsch et al., 2006; Harzsch, Wildt, Battelle, & Waloszek, 2005; Nässel, 1999; Schmid & Duncker, 1993; Sombke & Harzsch, 2015). A soma cortex including three discernable protocerebral clusters surrounds most of the central nervous system (CNS) (Figs. 2; 5a; 6a-b, d; 7a–d). Soma cluster 1 (SC1) is located between the first order visual neuropils of the lateral eyes and the second order visual neuropil of anterior lateral and posterior lateral eyes (AL2/PL2), cluster 2 (SC2) medially to the AL2/PL2, and cluster 3 (SC3) anteromedially to the mushroom body bridge (MBbr) (Fig. 2).

**Figure 1.**
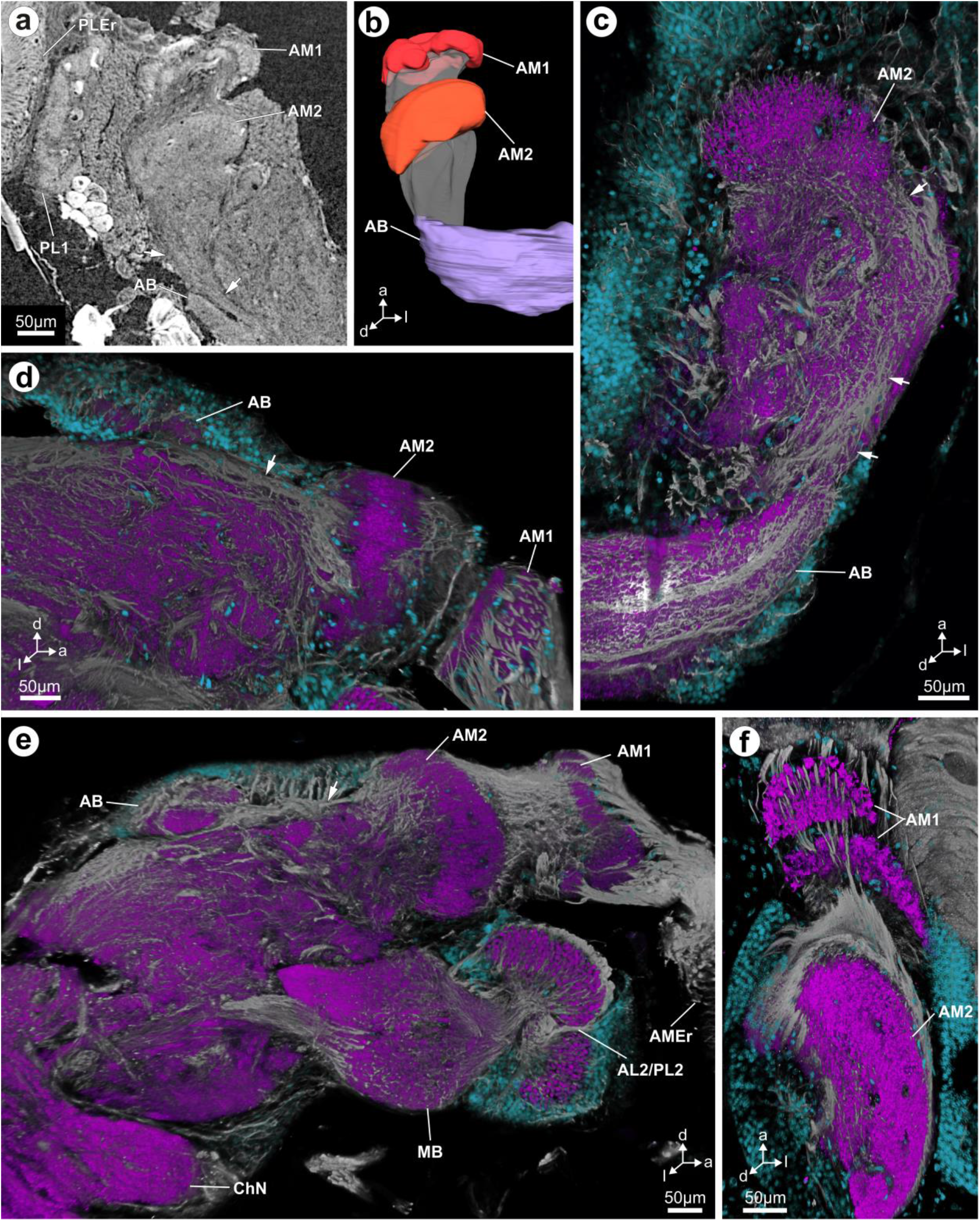
The principal eye pathway in *Marpissa muscosa*. **a** Virtual horizontal section of the dorsal protocerebrum obtained by microCT analysis shows the connections between the AM1, AM2 and the AB. **b** Three-dimensional reconstruction based on a microCT image stack shows the same connections as in a. **c-f** Maximum projection of image stacks (clsm), show tubulin-immunoreactivity (grey) and somata (blue) in the protocerebrum of *Marpissa muscosa*. **c** Horizontal section showing the AM2, the AB and connecting neurites (arrows). **D** Sagittal section through the AM1, AM2 and the AB showing the connecting neurites between AM2 and AB (arrow). **e** Sagittal section showing at least two lobes of the AM1, the thick tract to the AM2 and the thinner connection to the AB (arrow). AL2/PL2 and MB are also visible. **f** Horizontal section showing two lobes of the AM1, retinula cell axons terminating in those and interneurons connecting AM1 with AM2. **Abbreviations: a** anterior, **AB** arcuate body, **AL2/PL2** second order visual neuropil of anterior lateral and posterior lateral eyes, **AM1** first order visual neuropil of the anterior median eyes, **AM2** second order visual neuropil of the anterior median eyes, **AMEr** retina of the anterior median eyes, **ChN** cheliceral neuropil, **d** dorsal, **l** lateral, **MB** mushroom body, **PL1** first order visual neuropil of the posterior lateral eyes, **PLEr** retina of the posterior lateral eyes, **PM1** first order visual neuropil of the posterior median eyes.

**Figure 2.**
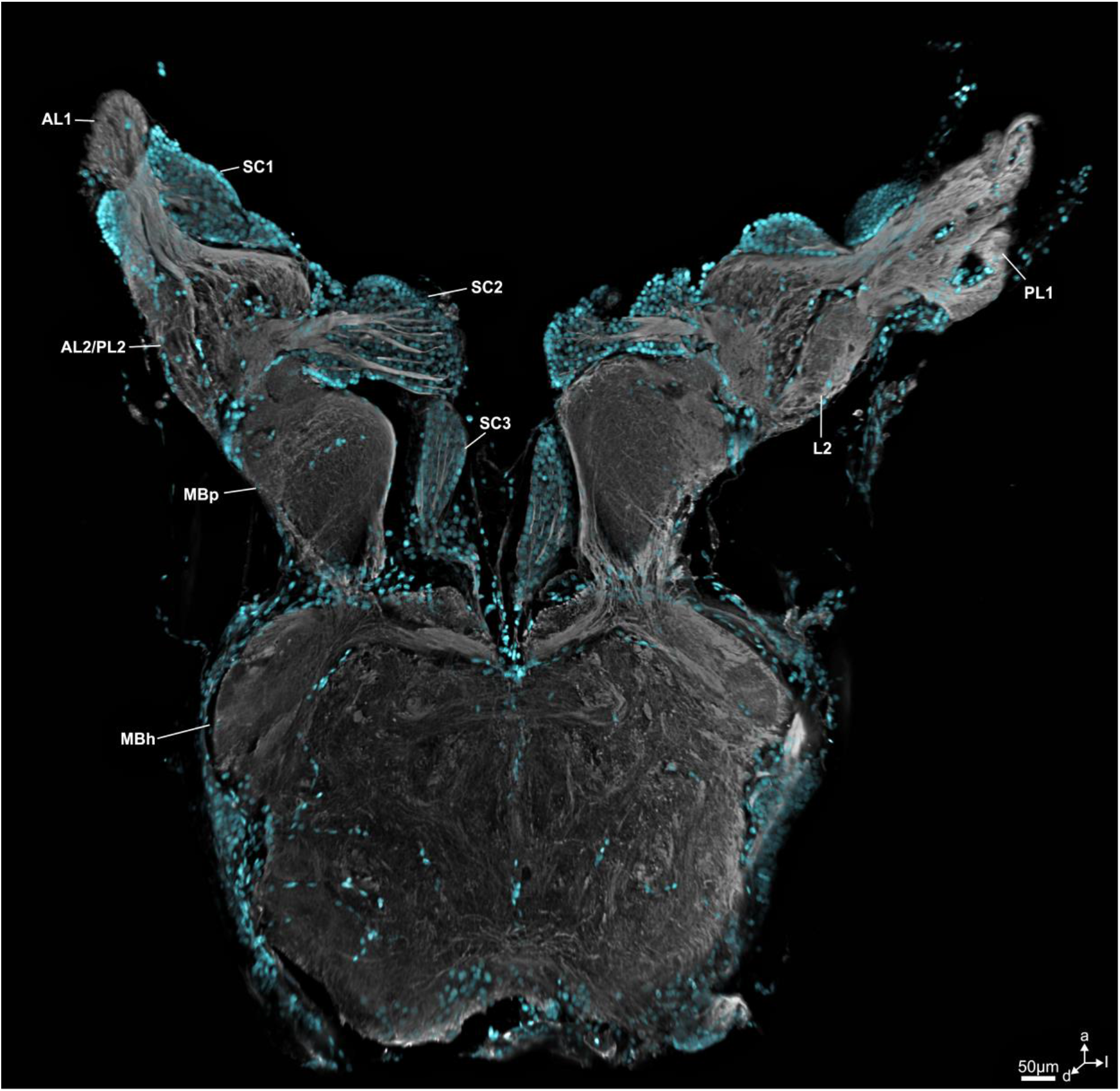
Maximum projection of an image stack (clsm), showing tubulin-immunoreactivity (grey) and somata (blue) in the protocerebrum of *Marpissa muscosa*. The AL1 and PL1 are connected to both, the AL2/PL2 and the L2. The neurites forming these connections originate in SC1. The AL2/PL2 and the L2 are connected to the MBp via neurites, which originate in the SC2. The two MBp are connected with each other via the mushroom body bridge, which is formed by long neurites that project along the periphery of the MBp and seem to originate in the SC2. Some of these long neurites also connect the MBp with the MBh. Other neurites originate in SC3 and enter the protocerebrum medially. **Abbreviations: a** anterior, **AL1** first order visual neuropil of the anterior lateral eyes, **AL2/PL2** second order visual neuropil of ALE and PLE, **d** dorsal, **l** lateral, **L2** shared second order visual neuropil of the anterior lateral and posterior lateral eyes, **MBh** mushroom body haft, **MBp** mushroom body pedunculus, **SC1-3** soma cluster 1-3.

**Figure 3.**
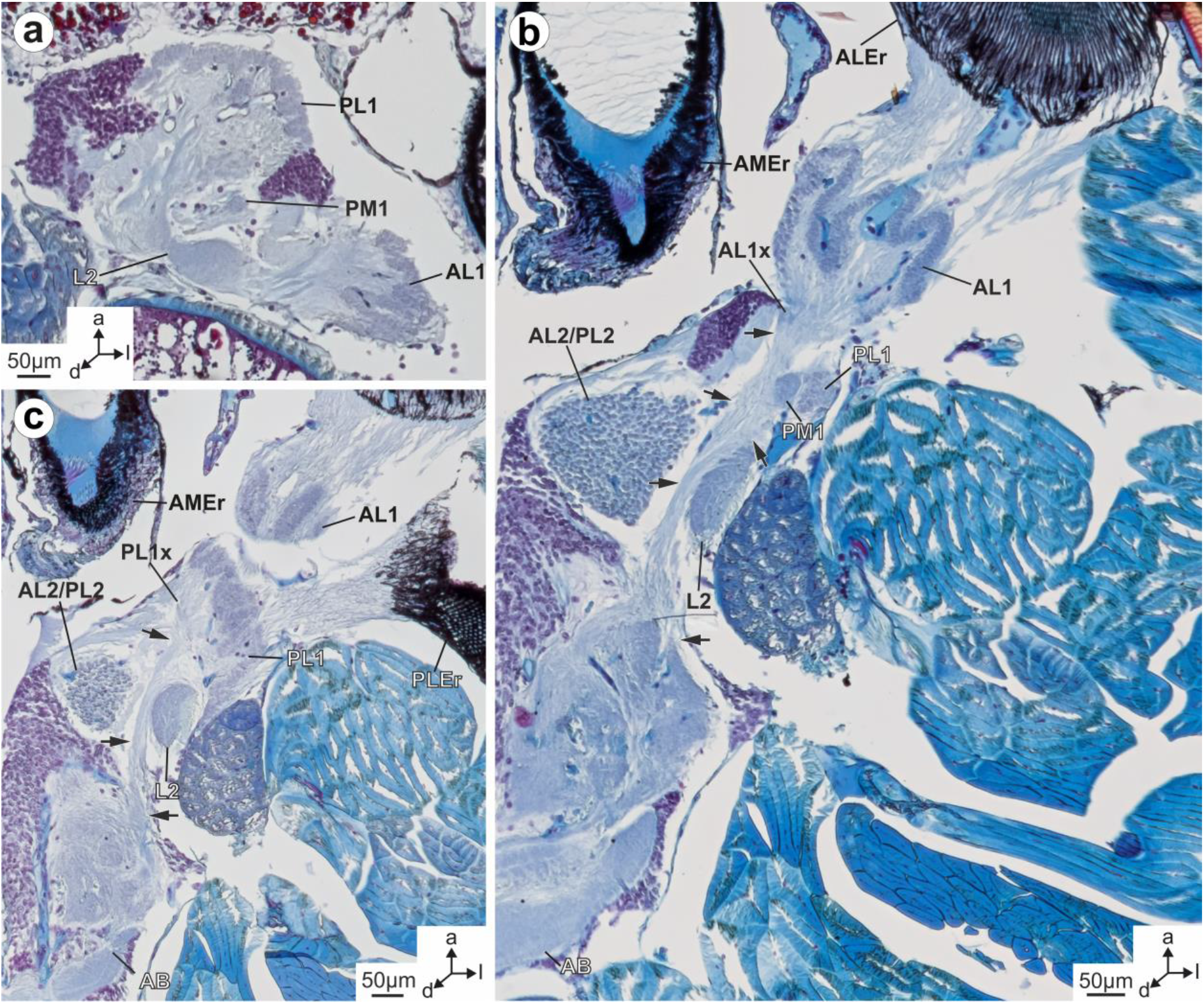
Paraffin sections through the anterior protocerebrum of *Marpissa muscosa.* **a** Sagittal section showing the conspicuous tracts that connect the AL1 and PL1 with the L2. **b** Horizontal section showing the convoluted and columnar structure of the AL1. Neurites that connect the AL1 with the AB probably synapse in the AL1x. Long neurites connecting the PL1 with the AB are visible curving around L2. Other long neurites contributing to the tract to the AB synapse in PM1. Arrows point to neurites contributing to the tract between first order visual neuropils of the secondary eyes and the arcuate body. **c** Horizontal section showing several neurites that connect the PL1 with the AL2/PL2, as well as the neurites contributing to the tract between the PL1 and the AB (highlighted by arrows). **Abbreviations: a** anterior, **AB** arcuate body, **AL1** first order visual neuropil of the anterior lateral eyes, **AL1x** neuropilar subunit of the AL1, **AL2/PL2** second order visual neuropil of anterior lateral and posterior lateral eyes, **ALEr** retina of the anterior lateral eyes, **AMEr** retina of the anterior median eyes, **d** dorsal, **l** lateral, **L2** shared second order visual neuropil of the anterior lateral and posterior lateral eyes, **PL1** first order visual neuropil of the posterior lateral eyes, **PLEr** retina of the anterior lateral eyes, **PL1x** neuropilar subunit of the PL1, **PM1** first order visual neuropil of the posterior median eyes.

**Figure 4.**
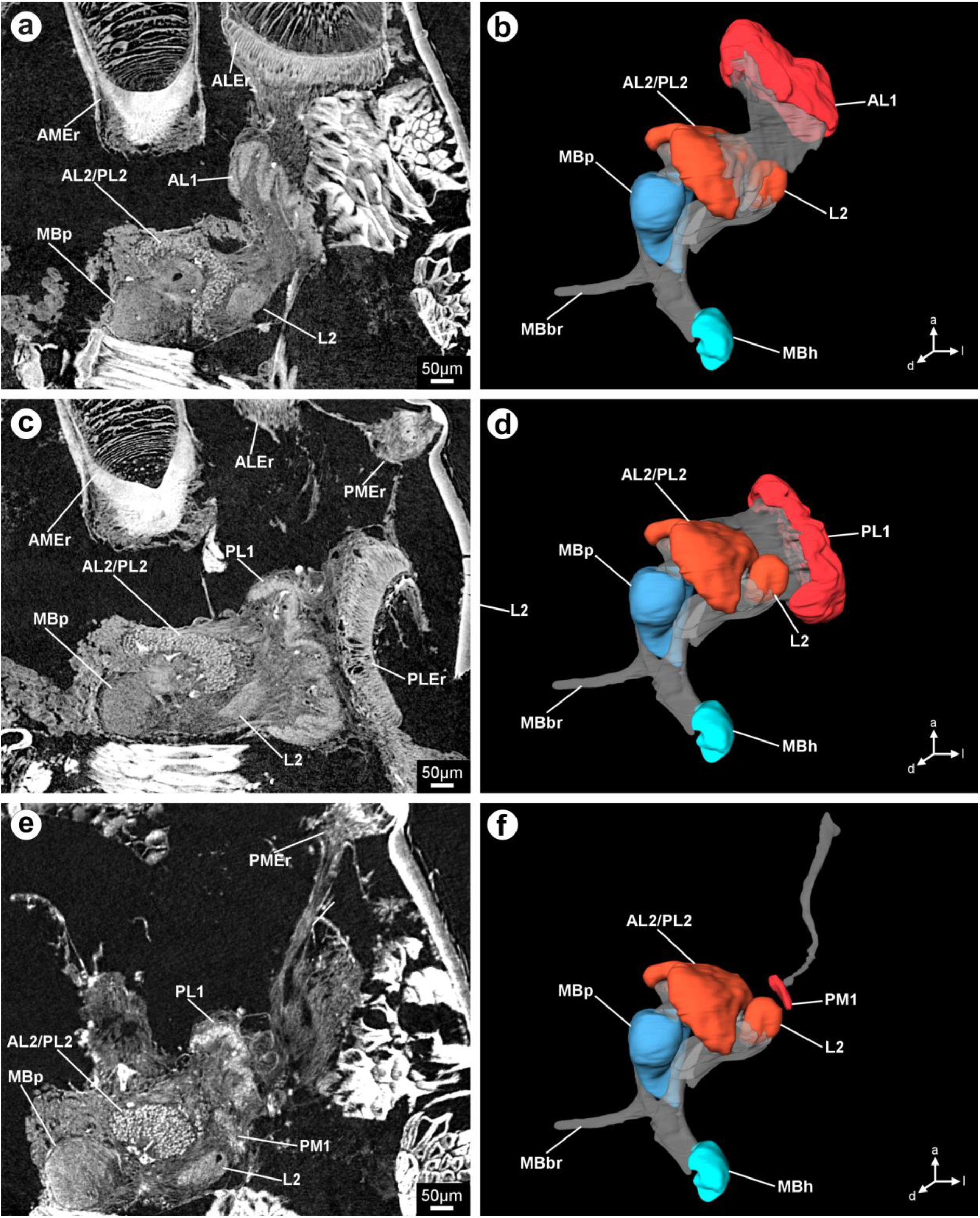
Protocerebral neuropils depicted in different virtual horizontal sections obtained by microCT analysis (**a, c, e**) and three-dimensional reconstructions based on microCT image stacks (**b, d, f**). **a** + **b** The retina of the ALE is located very closely to its first order visual neuropil, the AL1. The AL1 has a convoluted surface and is connected to the AL2/PL2 and the L2. **c** + **d** The retina of the PLE almost comes in contact with its first order visual neuropil, the PL1. The PL1 has a convoluted surface and is connected to the MB and the L2. **e**+ **f** A comparatively long optic nerve connects the retina of the PME with its first order visual neuropil, the PM1. See supporting information for an interactive 3D-visualization of the microCT reconstruction. **Abbreviations: a** anterior, **AL1** first order visual neuropil of the anterior lateral eyes, **AL2/PL2** second order visual neuropil of anterior lateral and posterior lateral eyes, **ALEr** retina of the anterior lateral eyes, **AMEr** retina of the anterior median eyes, **d** dorsal, **l** lateral, **L2** shared second order visual neuropil of the anterior lateral and posterior lateral eyes, **MBbr** mushroom body bridge, **MBh** mushroom body haft, **MBp** mushroom body pedunculus, **PL1** first order visual neuropil of the posterior lateral eyes, **PLEr** retina of the posterior lateral eyes, **PM1** first order visual neuropil of the posterior median eyes, **PMEr** retina of the posterior median eyes.

**Figure 5.**
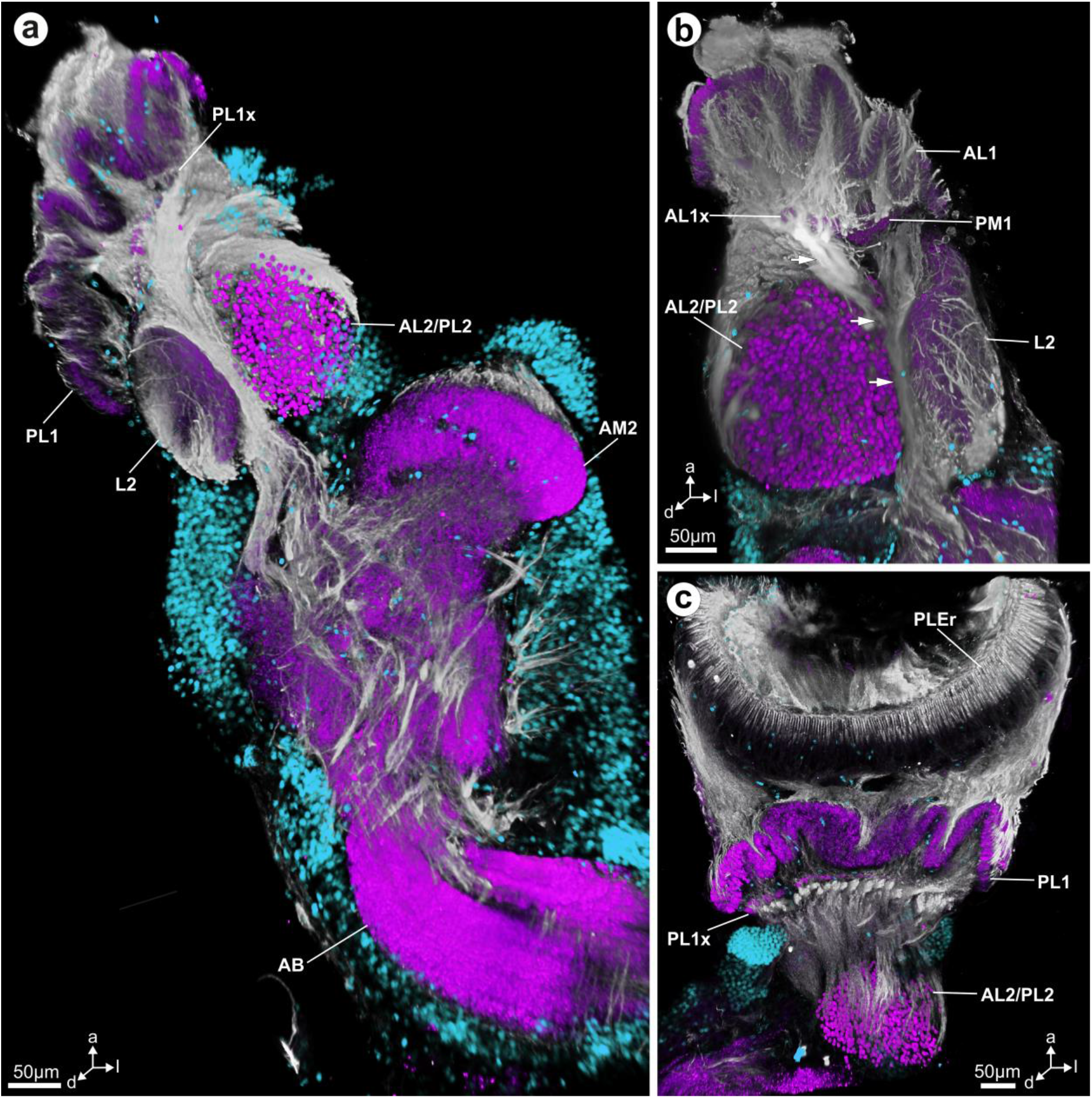
Maximum projections of image stacks (clsm) showing tubulin-immunoreactivity (grey), synapsin-immunoreactivity (magenta) and somata (blue) in the secondary eye visual pathway of *Marpissa muscosa*. **a** Neurites from the PL1 pass between AL2/PL2 and L2 towards the AB. **b** Neurites from AL1x join the tract from the PM1 that passes through between AL2/PL2 and L2 (highlighted by arrows). **c** The PL1 is in close spatial relationship to the PLEr. Parallel neurites connect the PL1 and PL1x with the Al2/PL2. **Abbreviations: a** anterior, **AB** arcuate body, **AL1** first order visual neuropil of the anterior lateral eyes, **AL1x** neuropilar subunit of the AL1, **AL2/PL2** second order visual neuropil of anterior lateral and posterior lateral eyes, **AM2** second order visual neuropil of the anterior median eyes, **d** dorsal, **l** lateral, **L2** shared second order visual neuropil of the anterior lateral and posterior lateral eyes, **MBp** mushroom body pedunculus, **PL1** first order visual neuropil of the posterior lateral eyes, **PL1x** neuropilar subunit of the PL1, **PLEr** retina of the posterior lateral eyes, **PM1** first order visual neuropil of the posterior median eyes.

**Figure 6.**
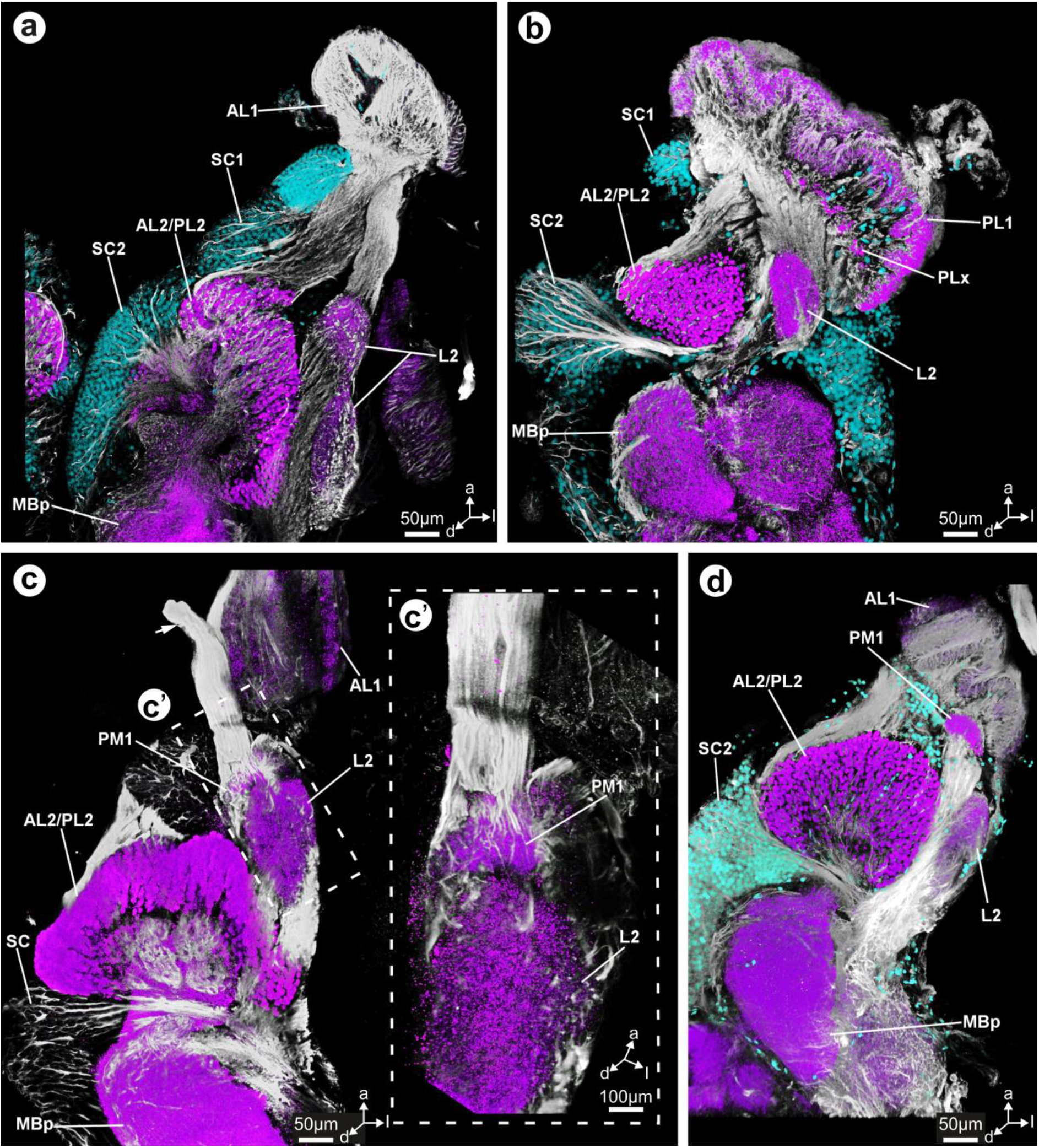
Maximum projections of image stacks (clsm) showing tubulin-immunoreactivity (grey), synapsin-immunoreactivity (magenta) and somata (blue) in the secondary eye visual pathway of *Marpissa muscosa*. **a** The AL1 is connected to the AL2/PL2 and the L2 via two prominent neurite tracts. Some of the neurites contributing to these tracts originate in SC1. Neurites connecting the AL2/PL2 with the MBp originate in SC2. **b** The PL1 is connected to the AL2/PL2 and the L2 via two prominent neurite tracts. Some neurites that arise in SC2 pass between AL2/PL2 and MBp towards the L2. **c** The optic nerve of the PME (highlighted by arrow) terminates in the PM1. **c’** Higher magnification of C, showing the termination site of the PME optic nerve in the PM1. **d** Neurites from the PM1 pass between AL2/PL2 and L2. **Abbreviations: a** anterior, **AL1** first order visual neuropil of the anterior lateral eyes, **AL2/PL2** second order visual neuropil of anterior lateral and posterior lateral eyes, **d** dorsal, **l** lateral, **L2** shared second order visual neuropil of the anterior lateral and posterior lateral eyes, **MBp** mushroom body pedunculus, **PL1** first order visual neuropil of the posterior lateral eyes, **PL1x** neuropilar subunit of the PL1, **PM1** first order visual neuropil of the posterior median eyes, **SC1-2** soma cluster 1-2.

**Figure 7.**
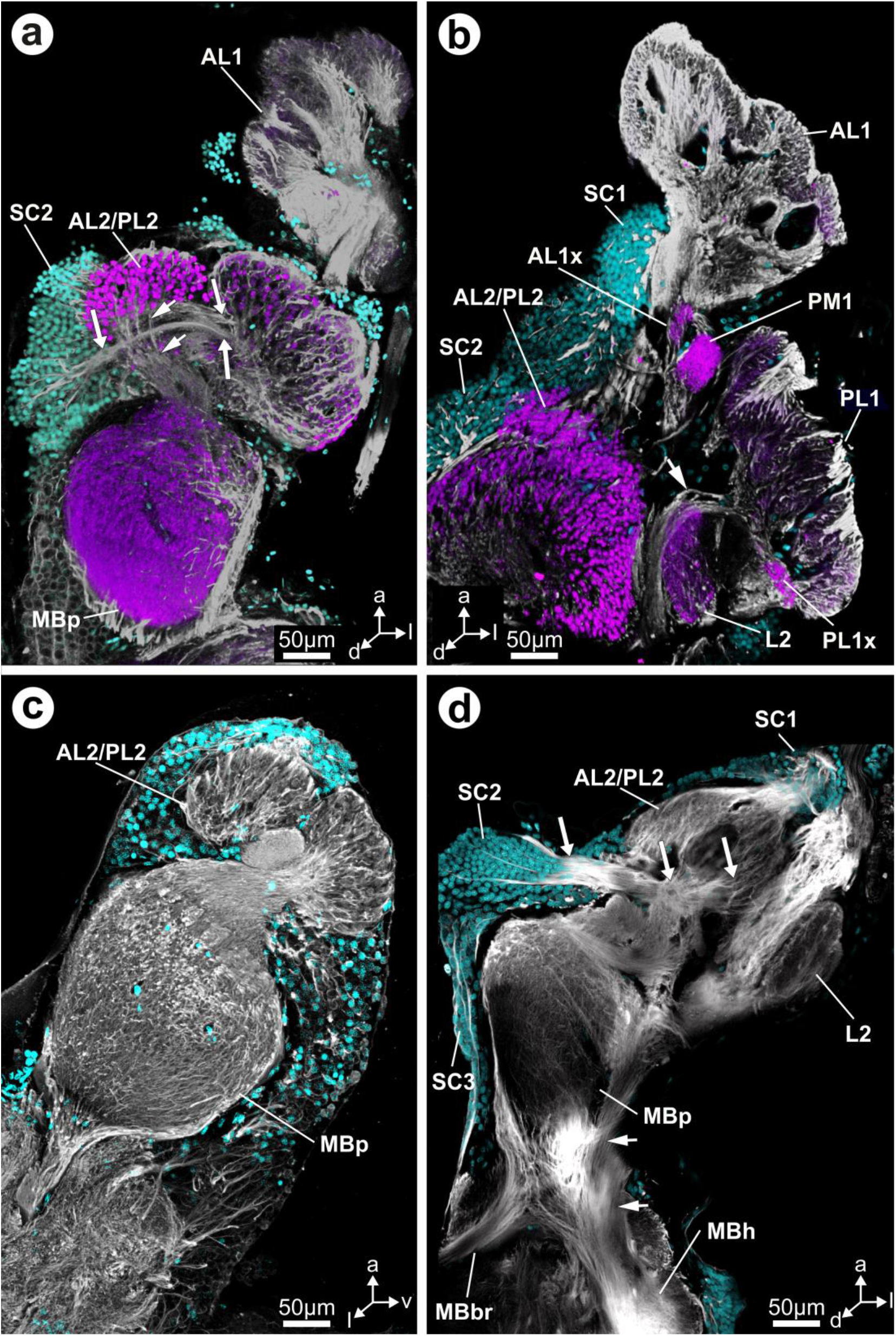
Maximum projections of image stacks (clsm) showing tubulin-immunoreactivity (grey), synapsin-immunoreactivity (magenta) and somata (blue) in the secondary eye visual pathway of *Marpissa muscosa*. **a** Horizontal section through AL1, MBp and AL2/PL2. The AL1 connects to one of the two tightly adjoining clusters of AL2/PL2. Arrows point to primary neurites connecting SC2 and AL2/PL2. Arrowheads point to neurites connecting the AL2/PL2 with the MBp. **b** Horizontal section through visual neuropils. The PM1 is situated between AL1 and PL1, AL2/PL2 appear as a uniform array. The arrowhead points to neurites connecting PL1 with AB, neurites probably synapse in the PL1x. **c** A sagittal section through AL2/PL2 and MBp reveals two distinct clusters of AL2/PL2 and the chiasm formed by neurites that connect AL2/PL2 with MBp. **d** Tubulin-immunoreactivity shows long peripheral neurites contributing to the MBbr and connecting the MBp with the MBh. Arrows point to primary neurites connecting SC2 with the AL2/PL2. Arrowheads point to neurites connecting the MBp with the MBh. **Abbreviations: a** anterior, **AL1** first order visual neuropil of the anterior lateral eyes, **AL1x** neuropilar subunit of the AL1, **AL2/PL2** second order visual neuropil of anterior lateral and posterior lateral eyes, **d** dorsal, **l** lateral, **L2** shared second order visual neuropil of the anterior lateral and posterior lateral eyes, **MBbr** mushroom body bridge, **MBh** mushroom body haft, **MBp** mushroom body pedunculus, **PL1** first order visual neuropil of the posterior lateral eyes, **PL1x** neuropilar subunit of the PL1, **PM1** first order visual neuropil of the posterior median eyes, **SC1-3** soma cluster 1-3.

**Figure 8.**
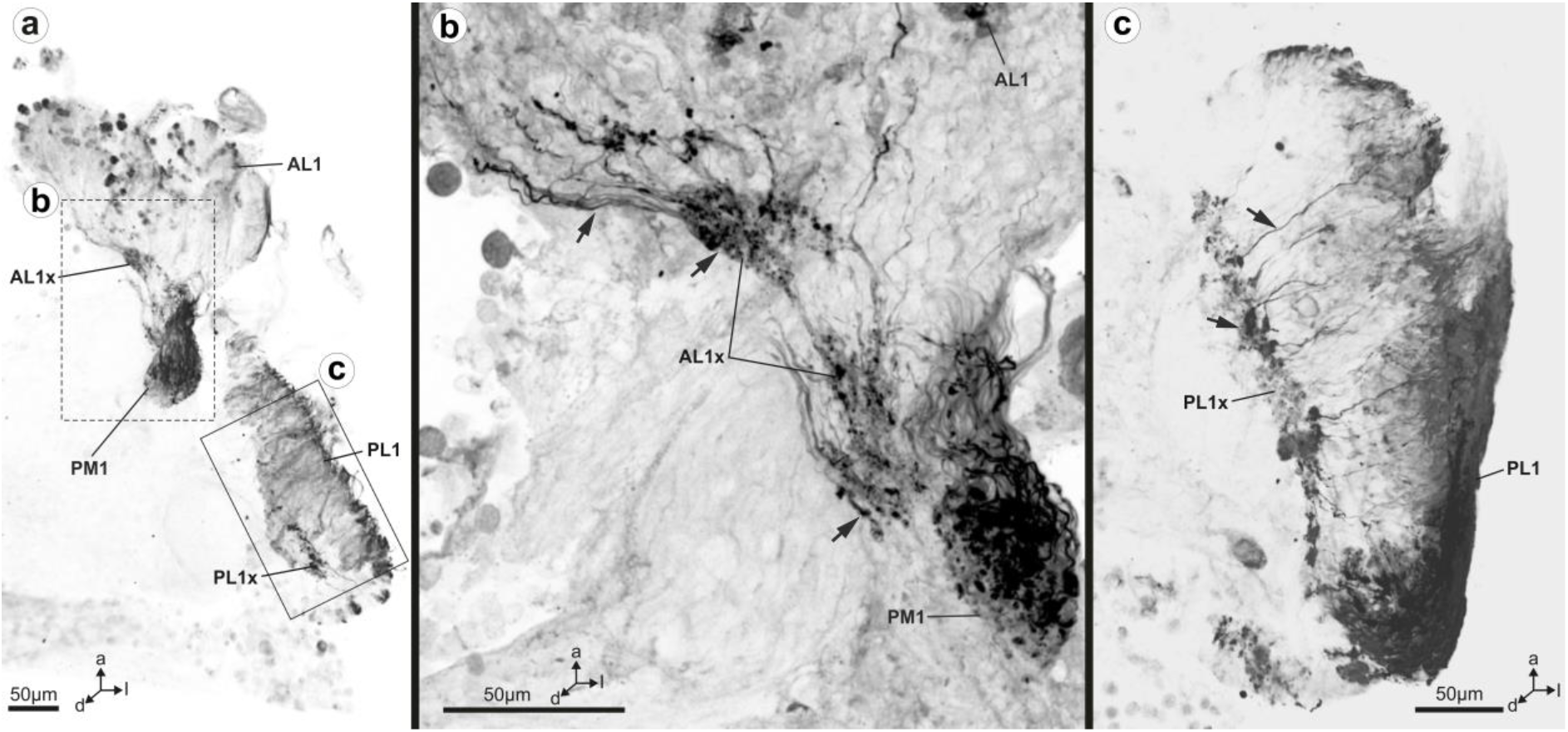
Maximum projections of image stacks (clsm), showing histamine-immunoreactivity in horizontal sections through the anteriormost protocerebrum of *Marpissa muscosa*. **a** Overview, shows strong histamine-immunoreactivity of retinula cell axon terminals in the AL1 and its substructure the AL1x, as well as in the PL1 and its substructure the PL1x. Retinula cell axons of the PME and their terminals in the PM1 exhibit strong histamine-immunoreactivity. **b** Magnification showing histamine-immunoreactive retinula cell axons and their terminals in the AL1x and PM1. Note the close spatial association of AL1x and PM1; AL1x is somewhat distorted due to sectioning. Arrows point to histamine-immunoreactive retinula cell axons and their terminals. **c** Magnification showing histamine-immunoreactive retinula cell axons and their terminals in the PL1 and PL1x. Arrows point to a retinula cell axon and a large terminal in the PL1x. **Abbreviations: a** anterior, **AL1** first order visual neuropil of the anterior lateral eyes, **AL1x** neuropilar subunit of the AL1, **d** dorsal, **l** lateral, **PL1** first order visual neuropil of the posterior lateral eyes, **PL1x** neuropilar subunit of the PL1, **PM1** first order visual neuropil of the posterior median eyes.

### 3.2 The principal eye pathway

The retina of the anterior median eyes (AME) is connected to its first order visual neuropil (AM1), which consists of at least two, but most likely four tightly adjoining subunits (Fig. 1a,e,f). The AM1 is situated close to the second order visual neuropil of the AME (AM2) and connected to it *via* parallel neurites (Fig. 1e,f). The AM2 is large, oval shaped and its posterior margin within the protocerebral neuropil is not clearly recognizable in anti-synapsin labeling (Fig. 1). Anti-tubulin labeling reveals a tract of neurites that connects the AM2 with the lateral flanges of the AB, as well as some neurites with unclear termination that penetrate the protocerebral neuropil (Fig. 1c-e).

### 3.3 Connectivity of the first order visual neuropils of the anterior and posterior lateral eyes

The retinae of the anterior lateral eyes (ALE) and posterior lateral eyes (PLE) are connected with their respective first order visual neuropils (AL1 and PL1) *via* short, thick optic nerves (Figs. 3b, c; 4a-b; 5c). A subpopulation of histaminergic neurites, which are interpreted here as retinula cell axons, terminates in distinct distal layers of these neuropils (AL1x and PL1x; Fig. 8a–c). Both the AL1 and PL1 are connected to the second order visual neuropil of the lateral eyes (L2) and the AL2/PL2 *via* prominent tracts of parallel ordered neurites that exhibit strong tubulin-immunoreactivity (Figs. 2; 5c; 6a-b, d; 7a-b). This connectivity is also evident in both, histological sections (Fig. 3) and microCT analysis (Fig. 4). The tracts are predominantly formed by neurites that originate in SC1, which surrounds this region of the anterior protocerebrum laterally, dorsally and medially (Figs. 2; 6a-b; 7b, d). The L2 and the AL2/PL2 are bi-lobed, which corresponds to the separate inputs from AL1 and PL1 (Figs. 3a; 3b, d; 5a; 6a, C; 8a, b). In histological sections, a thin but conspicuous tract is detectable that passes between L2 and AL2/PL2 and likely terminates in the arcuate body (AB) (Figs. 3b, c; 5a, b; 6d; 7b). The neurites forming this tract might originate in SC1 and further innervate the AL1x and PL1x, as well as the PM1 (see below; Figs. 3; 5a, b; 6d).

### 3.4 Connectivity of the first order visual neuropils of the posterior median eyes

The posterior median eyes (PME) are much smaller than all three other pairs of secondary eyes, and so is their retina (Fig. 4c, e). Because of the small retina, only a thin but long nerve, which enters the protocerebrum between the AL1 and PL1, connects the PME to its first order visual neuropil (PM1) (Figs. 4e; 6c, c’). The PM1 is situated between AL1 and PL1 (Figs. 3a-b; 4e, 5b; 6c-d; 7b; 8a). A prominent tract projects from the PM1 to the dorsoventral part of the protocerebrum and seems to terminate in the anterolateral apices of the AB (Fig. 3b; 5b; 6d).

### 3.5 The second order visual neuropil of ALE and PLE and the mushroom body

The second order visual neuropil of ALE and PLE (AL2/PL2) is of glomerular organization and composed of two lobes that tightly adjoin dorsally and are fused in their ventral part (Figs. 7a, c, d; 9a-b). Thus, only a uniform array of glomeruli is visible in most sections (Figs. 3b; 4a, c, e; 5; 6). Each glomerulus is composed of a high number of synaptic complexes (Fig. 9c–e). These synaptic complexes can be visualized by anti-synapsin labeling and phalloidin-labeling (Fig. 9d, e). Co-localization of synapsin and phalloidin is almost absent (Fig. 9). Individual glomeruli are innervated by neurites originating in the AL1 and PL1 (Figs. 5c; 6a, b, d; 7d). Anti-tubulin labeling reveals that neurites interweave the array of glomeruli mostly in parallel from proximal to distal (Fig. 5c; 6a). Within the core-region of the AL2/PL2, tubulin-immunoreactive neurites are more abundant (Fig. 6a,c. Two neurite bundles (MBn) connect the AL2/PL2 and the mushroom body pedunculus (MBp) and form a distinct chiasm close to the AL2/PL2 (Figs. 7a, c, d; 9a). Neurites from the medial SC2 contribute to these bundles by projecting into the AL2/PL2 and into the MBp (Figs. 2; 6a-c; 7a, d). Other neurites from SC2 pass between AL2/PL2 and MBp towards the L2 (Fig. 6b, d). Thus, a large number of interweaving neurites characterizes the region of the MBn, which makes tracing of individual trajectories difficult (Fig. 7a, d).

**Figure 9.**
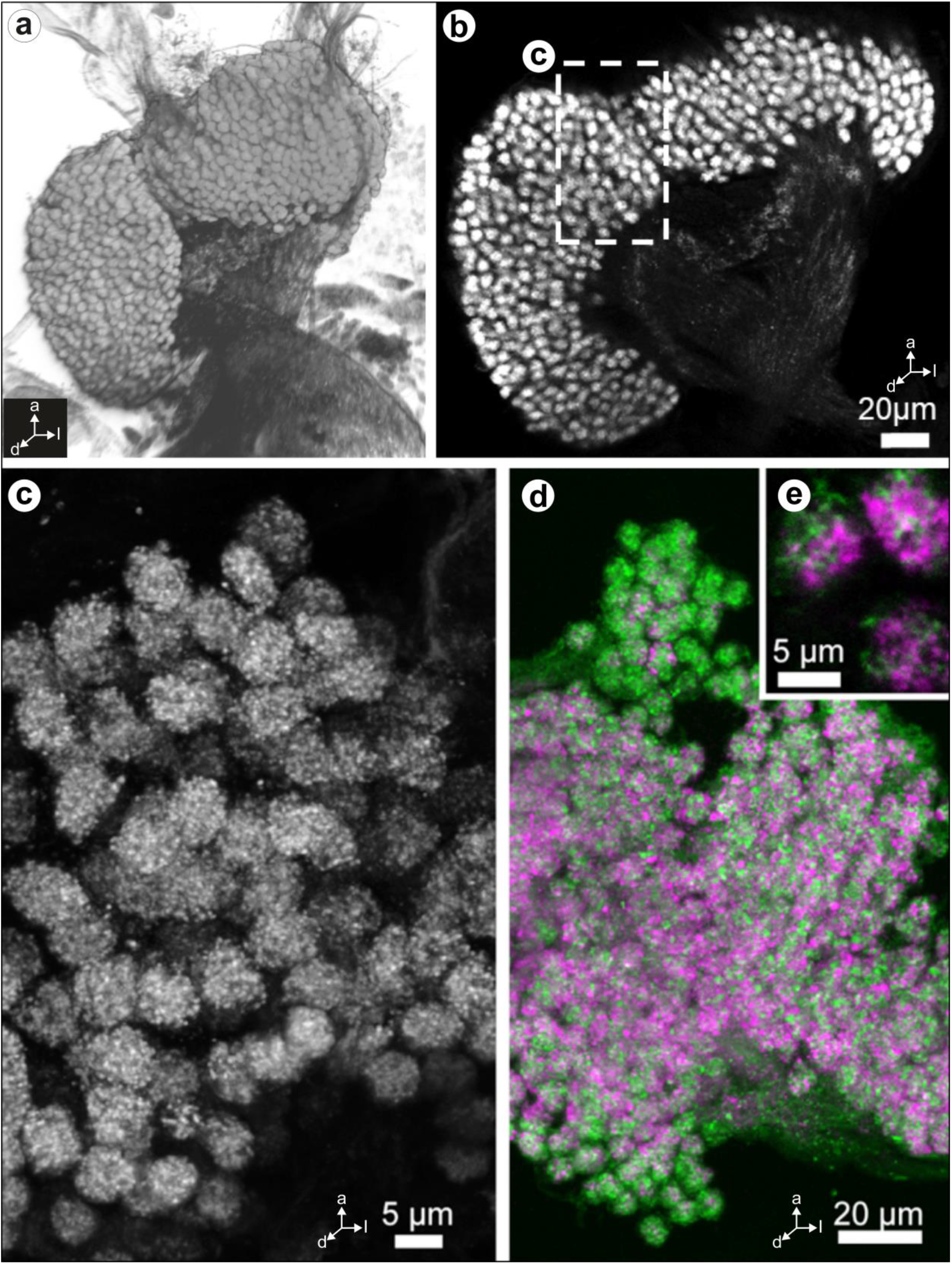
The second order visual neuropil of anterior lateral and posterior lateral eyes in the secondary eye visual pathway of *Marpissa muscosa.* **a** Three-dimensional volume rendering of an anti-synapsin labeled whole-mount, view from posterodorsal. Note the two distinct clusters of glomeruli that adjoin closely and coat the MBp anteriorly. **b** Single optical section of the AL2/PL2 within an anti-synapsin labeled whole-mount shows two distinct clusters of AL2/PL2 and a chiasm of neurites connecting them to the MBp. **c** Stack of optical sections through the AL2/PL2 (maximum intensity projection), showing synapsin-immunoreactivity. The glomerular subunits within each individual AL2/PL2 may represent synaptic boutons. **d** Anti-synapsin (magenta) and phalloidin (green) double–labeling reveals glomerular substructures (boutons) of the AL2/PL2. **e** Three individual AL2/PL2 with glomerular substructures (boutons) labeled by anti-synapsin and phalloidin.

Some long neurites that appear to originate in SC2 project along the periphery of the MBp and join with neurites arising from the mushroom body haft (MBh). Together they form the mushroom body bridge (MBbr) (Figs. 2; 6c; 7a, c, d). Other peripheral neurites from SC2 enter the MBh, where they seem to terminate (Figs. 2; 4b, d, f; 7d)

### 3.6 Visual neuropils and the mushroom body in *Cupiennius salei*

The dorsalmost neuropils in the protocerebrum of *C. salei* are the AM1 and AM2, which are connected by numerous intertwining neurites (Fig. 10a). The AM2 is connected to the AB *via* a thin tract (Fig. 10a). Located ventrally and somewhat anterior to the AM1 is the PM1, which is large and spans over one hemisphere of the protocerebrum at its broadest part (Fig. 10a,b,e). Its shape is cup-like, but somewhat irregular (Fig. 10e). The PL1 is situated anterioventral of the PM1, is similar in size, but horseshoe shaped (Fig. 10b). Both the PM1 and the PL1 are subdivided into an upper, strongly synapsin-immunoreactive part where the retinula cell axons terminate and a lower part where the synapsin-immunoreactivity is more patchily distributed (PM1x and PL1x; Fig. 10b,e). Tubulin-immunoreactivity is strongest in the lower part, where the neurites that connect the PM1 and PM2 with their respective second order neuropils bundle together. The AL1 is the smallest first-order neuropil and located close to the midline, between PM1 and PL1 (Fig. 10b). Each first order visual neuropil is connected to its second order visual neuropil. These are of glomerular organization and situated close to each other (Fig. 10b,c). While the AL2 is horseshoe shaped, the PL2 and PM2 are more elongate in shape and also larger (Fig. 10b,c).

**Figure 10.**
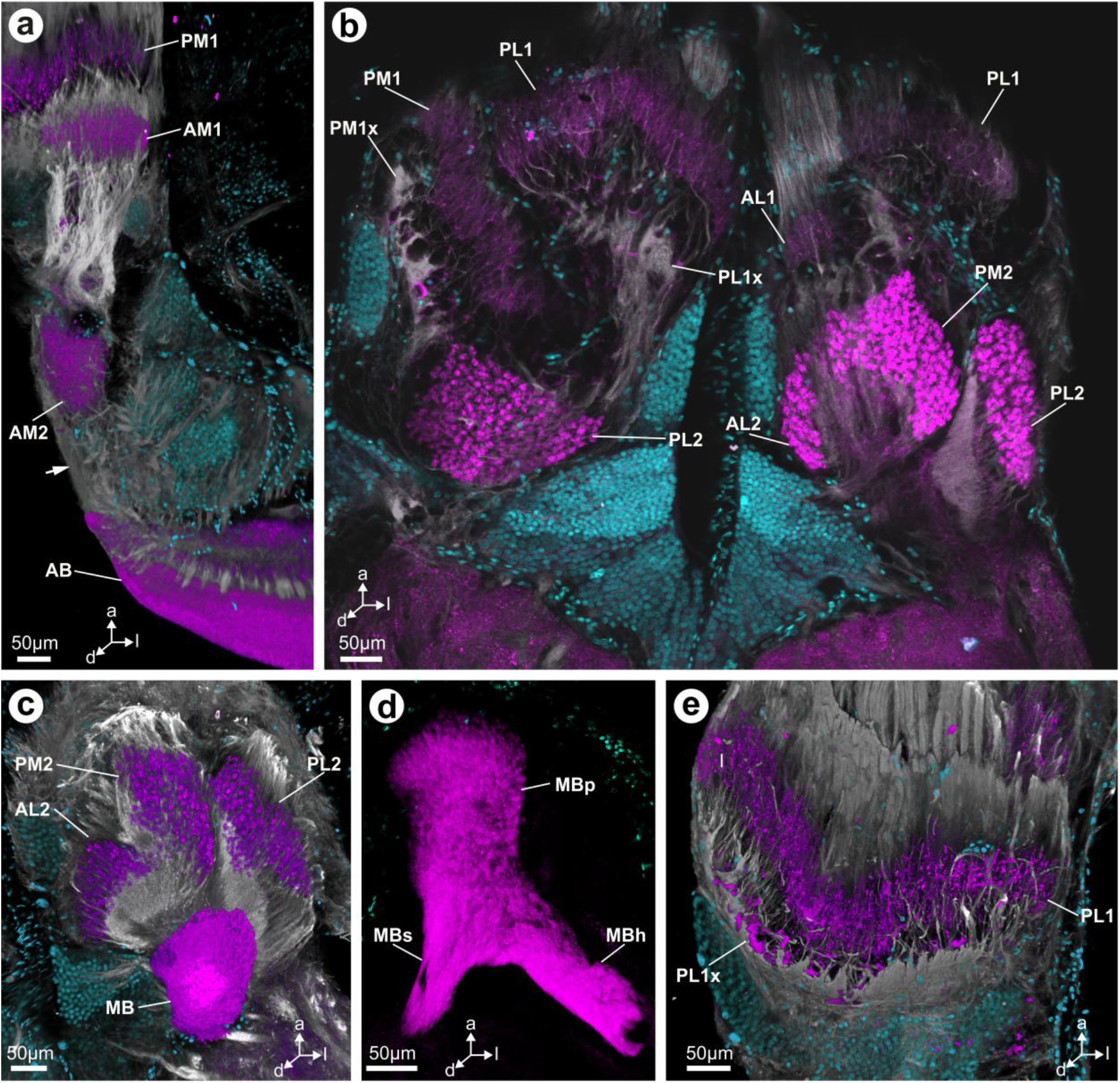
Maximum projections of image stacks (clsm; **a,c-e**) and single optical section (clsm) showing synapsin-immunoreactivity (magenta), tubulin-immunoreactivity (grey) and somata (cyan) in the secondary eye visual system of *Cupiennius salei*. **a** In the principal eye visual pathway, the AM1 is connected to the AM2 via a thick tract. Connecting neurites between AM2 and the AB are also visible (arrow). **b** Within the secondary eye visual pathways, every eye is associated with its own first order visual neuropil (AL1, PM1 and PL2); the PM1 and PL1 appear to have a substructure (PM1x and PL1x). All three second order visual neuropils are sub-structured into glomeruli. **c** The AL2, PM2 and PL2 are all connected to the MB. **d** The MB of *C. salei* consists of a MBp, MBh and MBs. All three substructures form a continuous synapsin-rich neuropil. **e** Horizontal section showing the PM1 and the PM1x. **Abbreviations: a** anterior, **AB** arcuate body, **AL1** first order visual neuropil of the anterior lateral eyes, **AL2** second order visual neuropil of the anterior lateral eyes, **AM1** first order visual neuropil of the anterior median eyes, **AM2** second order visual neuropil of the anterior median eyes, **d** dorsal, **l** lateral, **MBh** mushroom body haft, **MBp** mushroom body pedunculus, **MBs** mushroom body shaft, **PL1** first order visual neuropil of the posterior lateral eyes, **PL1x** neuropilar subunit of the PL1, **PL2** second order visual neuropil of the posterior lateral eyes, **PM1** first order visual neuropil of the posterior median eyes, **PM1x** neuropilar subunit of the PM1, **PM2** second order visual neuropil of the posterior median eyes.

The mushroom body is characterized by strong synapsin-immunoreactivity and consists of a pedunculus (MBp), a shaft (MBs) and a haft (MBh) (Fig. 10b). The MBs and the MBh are fused with the MBp, and together form one continuous synapsin-immunoreactive domain (Fig. 10b).

## 4 Discussion

Comparative neuroanatomical studies on different spider species show that while general brain anatomy is similar, the structure of different neuropils especially within the protocerebrum varies (Hanström, 1921; Long, 2016; Steinhoff et al., 2017; Weltzien and Barth, 1991). The previous investigation by Steinhoff et al. (2017) on the general brain anatomy of *Marpissa muscosa* suggested that the brains of jumping spiders and the spider species *Cupiennius salei* differ in structure and arrangement of neuropils. Although the general architecture is similar, we have shown here that the connectivity of neuropils in the secondary eye pathways differ between these two species, which suggests differences in information processing and function of secondary eye visual neuropils.

The relative size and position of the secondary eyes differs between *M. muscosa* and *C. salei.* While in *M. muscosa* the PME is very small and oriented sideways, it is the largest eye in *C. salei* and oriented forward (Morehouse et al., 2017; Fig. 11). The PLE are similar in relative size and position, but the ALE are considerably smaller in *C. salei,* albeit also oriented forward (Fig. 11). The relative size of the retinae as well as the brain is smaller in *C. salei*, which is why the optic nerves associated with the eyes are much longer in *C. salei* (Fig. 11).

**Figure 11.**
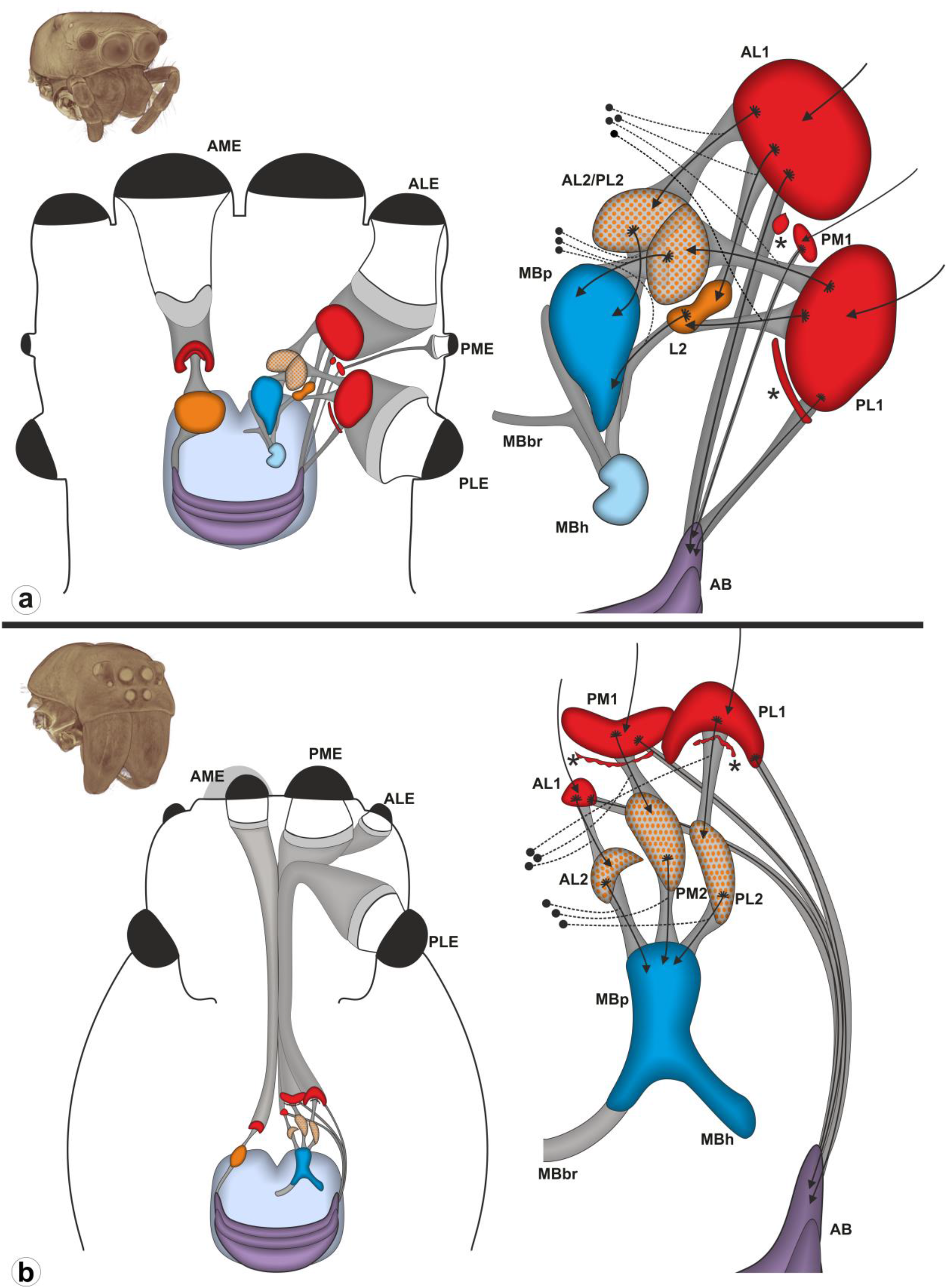
Schematic representations of the principal and secondary eye visual pathways in the brains of **a***Marpissa muscosa* and **b***Cupiennius salei.* Insets top left: 3D volume rendering of the prosomata of **a***M. muscosa* and **b***C. salei*, based on microCT analysis. **b** Left anterior median eye concealed by posterior median eye. **Abbreviations: AB** arcuate body, **AL1** first order visual neuropil of the anterior lateral eyes, **AL2** second order visual neuropil of the anterior lateral eyes, **AL2/PL2** second order visual neuropil of anterior lateral and posterior lateral eyes, **ALE** anterior lateral eye, **AME** anterior median eye, **L2** shared second order visual neuropil of the anterior lateral and posterior lateral eyes, **MBbr** mushroom body bridge, **MBh** mushroom body haft, **MBp** mushroom body pedunculus, **PL1** first order visual neuropil of the posterior lateral eyes, **PL2** second order visual neuropil of the posterior lateral eyes, **PLE** posterior lateral eye, **PME** posterior median eye, **PM1** first order visual neuropil of the posterior median eyes, **PM2** second order visual neuropil of the posterior median eyes. **Filled black circles** indicate somata with known positions, **dotted lines** represent primary neurites and **arrows** indicate the assumed direction of information flow. **Asterisks** mark neuropilar subregions of first order neuropils (AL1x and PL1x in *M. muscosa*, PM1x and PL1x in *C. salei*).

### 4.1 Connectivity of neuropils in the principal eye pathway

The connection of the AME retinula cell axons to the AM1 has recently been described in detail by Nagata et al. (2019) for the jumping spider *Hasarius adansoni*. These authors revealed that there are four photoreceptor layers in the retina, which each connect to one of four subregions in the AM1 (Nagata et al., 2019). We show here, that *in M. muscosa* the AM1 is further connected to the AM2, which is the same as described for *C. salei* (Strausfeld et al., 1993). In *C. salei*, two types of neurons have been identified whose axons connect the AM2 with the flanges and the inner part of the AB (Strausfeld et al., 1993). In *M. muscosa* this pattern is likely the same, although we were only able to unambiguously detect neurites that connect the AM2 with the flanges of the AB. There are, however, neurites that project from the AM2 into the protocerebral neuropil and some of those may well proceed towards the inner part of the AB.

### 4.2 Connectivity of the first order visual neuropils of the anterior and posterior lateral eyes

In *M. muscosa*, all retinula cell axons from ALE, PLE and PME seem to terminate in their associated first order visual neuropils (Fig. 8). This is in accordance to what has been found in other Araneae (Becherer and Schmid, 1999; Duelli, 1980; Schmid and Duncker, 1993; Strausfeld et al., 1993; Strausfeld and Barth, 1993). Nevertheless, this pattern is different in all other investigated taxa of Chelicerata (e.g. Scorpiones, Xiphosura, or Pycnogonida), where a subset of retinula cell axons also projects into the second order visual neuropils (Battelle, Sombke, & Harzsch, 2016; Lehmann & Melzer, 2013, 2018). It should be noted, however, that this discrepancy between spiders and all other arachnids still needs to be scrutinized using cobalt fills, which have not been employed in spiders yet (Lehmann and Melzer, 2018). In *C. salei*, a few histaminergic neurites terminate in all second order visual neuropils, and have been interpreted as interneurons (Schmid and Duncker, 1993). However, as long as no neuronal tracings have been carried out, it cannot be ruled out that these neurites might instead represent retinula cell axons. In *M. muscosa*, we did not detect any histaminergic neurites that terminate in the second order visual neuropils. There are, however, numerous histaminergic neurites that terminate in AL1x and PL1x and we interpret those as retinula cell axons.

In our histological sections, it appears as if some neurites that contribute to the tract between AL1, PL1 and the AB, arise in AL1x and PL1x respectively (Fig. 3b, c; 5a-b). Hill (1975) described in *Phidippus sp.* that neurites from the AL1x domain contribute to a tract projecting towards the posterior protocerebrum. However, Hill (1975) did not mention terminations of retinula cell axons in the AL1x and PL1x, but detected neurites connecting the AL1 with the AL1x. Thus, it is possible that Hill (1975) either misinterpreted retinula cell axons as interneurons, or that we missed connecting neurites between AL1 and AL1x. In our immunohistological investigations, we found that synapsin-immunoreactive substructures of first order visual neuropils are also present in *C. salei*, the PM1x and the PL1x. In both, *M. muscosa* and *C. salei,* this region is additionally characterized by strong tubulin-immunoreactivity. This can be explained by the tight packing of neurites that converge at the level of the AL1x and PL1x in *M. muscosa* and the PM1x and PL1x in *C. salei* to form the tracts to the respective second order visual neuropils. A subdivision of PM1 and PL1 is also visible in histamine-immunolabeled material from the study by Schmid & Duncker (1993; compare Figs. 2c,d and 3a,b), although not discussed by the authors. It remains open whether retinula cell axons or interneurons terminate at the synapsin immunoreactive part of the PL1x and PM1x in *C. salei* and the AL1x and PL1x in *M. muscosa*.

In *M. muscosa*, the convoluted first order visual neuropils of the ALE and the PLE are both connected with the AL2/PL2 and the L2, as well as with the AB, which is only partially consistent with Hill’s (1975) study on the nervous system in *Phidippus* spp., where he found that axons of the AL1, and not the PL1, join the tract that projects towards the AB. Duelli (1980), did not detect connections between PL1 and L2 in the salticids *Evarcha arcuta* (Clerck, 1757) and *Pseudamycus validus* (Thorell, 1877). He described “large field horizontal neurons,” closely bypassing the AL2/PL2 and L2 and connecting the PL1 with the posterior protocerebrum (Duelli, 1980). Judging from their structure and position, these neurons might be identical with the neurites described in our study which contribute to the tract that connects the first order visual neuropils of the secondary eyes with the arcuate body. This discrepancy might be due to incomplete staining in the earlier studies where reduced-silver staining was employed. Our approach combines specific and broader markers with specific staining techniques. It should also be noted that the studies on the secondary eye pathways in jumping spiders were conducted on different species. However, it seems unlikely that the secondary eye pathways differ considerably within salticoid jumping spiders (Maddison and Hedin, 2003; Morehouse et al., 2017).

Based on our results, AL2/PL2 and L2 are shared second order visual neuropils of both AL1 and PL1 in *M. muscosa*. The situation in *C. salei* is, however, different. Here, the first order visual neuropils of all three secondary eyes are connected to individual second order visual neuropils (compare Fig. 11a and b) (Strausfeld and Barth, 1993). Each of these second order visual neuropils (AL2, PL2, PM2) is composed of small neuropilar glomeruli (compare Fig. 11). An unstructured second order visual neuropil (without glomeruli), such as the L2, is absent in *C. salei* (Strausfeld and Barth, 1993). Similar to the “giant interneurons” described for jumping spiders by Duelli (1980), Strausfeld & Barth (1993) detected “lamina wide-field neurons” that connect all first order visual neuropils with a region close to the AB. It is very likely that these neurons correspond to the axons forming the tract between first order visual neuropils and the AB that we described in *M. muscosa*.

### 4.3 First order visual neuropils of the posterior median eyes

The PM1 is by far the smallest first order visual neuropil of all secondary eyes in *M. muscosa*. This correlates with the much smaller overall size of the PME (compared to ALE and PLE). The PME have previously been considered vestigial (Land, 1985a, 1972), but since there is an optic nerve and a corresponding first order visual neuropil, we conclude that these eyes are functional. Previously, Steinhoff et al. (2017) considered the PM1 to be a second order visual neuropil (“PM2”), based on its close spatial relationship with the AL1x. However, we now demonstrate that the PM1 is innervated only by retinula cell axons originating in the PME, and connections between PM1 and AL1x seem to be absent. This coincides with Hill’s (1975) description for *Phidippus* spp., although he described additional neurites connecting PL1 and PM1. It is likely that these neurites contribute to the tract that connects the first order visual neuropils with the AB (compare Fig. 11a), instead of actually forming synapses in the PM1. The PM1 differs from all other eyes by being connected to the AB only. Consequently, the pathway of the PME lacks a second order visual neuropil. Information from this eye is not processed within the mushroom bodies, but exclusively in the arcuate body.

### 4.4 Second order visual neuropils of the ALE and PLE

Visual neuropils with a glomerular structure have been found in the protocerebrum of several spider species (e.g. Long, 2016). They have been termed “lame glomérulée” by Saint-Remy (1887), “glomeruli” by Hanström (1921) and Babu & Barth (1984), “spherules” by Strausfeld & Barth (1993) and “microglomeruli” by Steinhoff et al. (2017). The presence and arrangement of these glomerular neuropils differs between spider species (Hanström, 1921; Long, 2016) and some authors have considered them as a substructure of the MB (Babu and Barth, 1984; Hill, 1975), whereas others have considered them to be separate second order visual neuropils (Long, 2016; discussed in Steinhoff et al., 2017; Strausfeld and Barth, 1993). Our results demonstrate, that in *M. muscosa*, the AL2/PL2 represents two closely adjoining second order visual neuropils of the ALE and PLE, respectively. In *M. muscosa*, the AL2/PL2 was also found to react less plastically to different environmental conditions than the MBp (Steinhoff et al., 2018), which was interpreted as further evidence for their function as discrete visual neuropils. Within the AL2/PL2, each glomerulus seems to consist of numerous synaptic complexes and Duelli (1980) estimated roughly 400 synapses per glomerulus in the jumping spider *Evarcha arcuta*. Studies in insects used anti-synapsin labeling to detect presynaptic sites, and phalloidin (which labels f-actin) to highlight postsynaptic sites of synaptic complexes (Groh and Rössler, 2011). To investigate the AL2/PL2 of *M. muscosa*, we have employed the same double-labeling approach (Fig. 9d-e), but the anti-synapsin and phalloidin labeled areas in each synaptic complex were not clearly differentiated, as is the case in social hymenopterans (Groh and Rössler, 2011), despite co-localization being minimal. This might indicate that phalloidin does not consistently label postsynaptic sites in *M. muscosa*, or that the organization of the pre- and postsynaptic sites within the AL2/PL2 of *M. muscosa* differs compared to the glomeruli in mushroom body calyces of investigated insect species. To clarify synaptic connectivity within the AL2/PL2 in *M. muscosa* and spiders in general, a detailed transmission electron microscopic study is needed.

### 4.5 Mushroom body pedunculus, haft and bridge

The somata within SC2 of the neurons that connect the second order visual neuropils to the MBp have been referred to as “globuli cells” (Hanström, 1921; Steinhoff et al., 2017) and the neurons including these somata as “T-cells” (Strausfeld and Barth, 1993). In *C. salei*, globuli cells have a smaller diameter of the soma and nucleus than other cell bodies in the protocerebrum (Babu and Barth, 1984). We could not find such size differences in *M. muscosa*. In fact, all three nuclear markers revealed that somata of the three identified clusters as well the somata of the surrounding cortex exhibit a similar size (Figs. 2; 3; 5; 6; 7). Thus, according to their definition (Kenyon, 1896; Richter et al., 2010), somata of SC2 are no globuli cells in *M. muscosa*. The long neurites that connect the SC2 with the AL2/PL2 and the L2 are probably primary neurites (Richter et al., 2010), from which dendrites branch into the AL2/PL2 and L2 respectively, and axons project into the MBp, MBh and MBbr. A similar arrangement has been described by Babu and Barth (1984) for *C. salei* and by Hill (1975) for two *Phidippus* species.

The structure and organization of the MB differs between *M. muscosa* and *C. salei*, as already reported by Steinhoff et al. (2017). The MBh is a separate structure of the mushroom body in *M. muscosa,* connected to the MBp by neurites (Steinhoff et al., 2017), whereas in *C. salei*, it is fused with the MBp (Strausfeld and Barth, 1993). Furthermore, the mushroom body shaft, which in *C. salei* gives rise to the MBbr (Strausfeld and Barth, 1993) is absent or fused with the MBp in *M. muscosa.*

### 4.5 Functional considerations

The jumping spider *Marpissa muscosa* and the wandering spider *Cupiennius salei* differ strongly in their lifestyle. While *M. muscosa* is a diurnal hunter that actively pursues its prey and relies on vision for prey capture, *C. salei* is an exclusively nocturnal sit-and-wait predator, which detects its prey based on vibratory cues (Barth & Seyfarth, 1979). Although attack behavior of *C. salei* can be elicited by visual cues under experimental conditions (Fenk et al., 2010), mechanosensing is the major modality for hunting prey, since *C. salei* also captures prey when its eyes are occluded (Hergenröder and Barth, 1983). Given these obvious differences in lifestyle and relative importance of sensory modalities, it is surprising how similar the general brain anatomy of *M. muscosa* and *C. salei* is. Apart from differences in size and structure of neuropils, the visual pathway of the principal eyes is identical in *C. salei* and *M. muscosa*. Parts of the visual pathways of the secondary eyes in the jumping spider *Marpissa muscosa* differ, however, from those in the wandering spider *Cupiennius salei.* These differences might reflect the differences in the visual fields and corresponding functions of their secondary eyes (Land, 1985b; Land and Barth, 1992). The most conspicuous difference between the visual systems in the brains of *M. muscosa* and *C. salei* is the pathway of the PME. While the two larger secondary eyes (ALE and PLE) are connected to the mushroom bodies in *M. muscosa,* the PME are not (Fig. 11). This suggests that the PME do not play a role in motion detection (Duelli, 1978; Land, 1971; Zurek et al., 2010). In *C. salei,* however, the PME are important movement detectors (Schmid, 1998) connected to the mushroom bodies (Strausfeld and Barth, 1993).

Another interesting difference between the two species is the presence of the L2 in *M. muscosa* (Steinhoff et al., 2017), which is absent in *C. salei*. The structure of this neuropil differs from the AL2/PL2 and all second order visual neuropils in *C. salei,* since the L2 in *M. muscosa* lacks a glomerular organization. Glomeruli might ensure a retinotopic organization of neurites within the second order visual neuropils. This has been hypothesized for both *C. salei* (Strausfeld and Barth, 1993) and for the PLE pathway of jumping spiders (Duelli, 1980), where no lateral connections between individual glomeruli of the AL2/PL2 were found. We hypothesize that while a retinotopic organization is kept in the AL2/PL2, it is lost in the L2. Information from the movement-detecting ALE and PLE in *M. muscosa* might be integrated not only in the MB, but also in the L2, and enable faster movement decisions. Alternatively, a glomerular organization might be the most space-saving structure for the processing of input from differentially tuned groups of axons (Eisthen, 2002; Hildebrand and Shepherd, 1997), and although the L2 is not composed of glomeruli, information might still be processed in parallel without lateral connections. This would also indicate that a higher number of neurites synapse in the AL2/PL2 than in the L2 in *M. muscosa*.

Some early-branched off salticid species possess a large PME that has its own field of view (e.g. *Portia*; Land, 1985b; Maddison and Hedin, 2003). Land (1985b) suggested that in jumping spiders with a small PME, such as *M. muscosa*, the fields of view of ALE and PLE have expanded to fill the space not covered by the PME. The evolutionary transformation to a smaller PME and larger ALE and PLE might have led to visual pathways that also incorporated the L2. Thus in early-branched off jumping spiders with large PME, it could be hypothesized that the L2 might serve as a second order visual neuropil of the PME. Future studies should thus analyze the brain architecture of early-branched off salticid species.

## AUTHOR CONTRIBUTIONS

All authors had full access to all the data in the study and take responsibility for the integrity of the data and the accuracy of data analysis. Study concept and design: POMS, AS, GU. Acquisition of data: POMS. Analysis and interpretation of data: POMS, AS. Wrote the manuscript: POMS, AS. Contributed to the writing of the manuscript: GU, SH. Obtained funding: GU, SH, POMS.

## ACKNOWLEDGEMENTS

The authors wish to thank Georg Brenneis and Matthes Kenning (University of Greifswald) as well as Nicholas J. Strausfeld (University of Arizona, Tucson) and Friedrich G. Barth (University of Vienna) for helpful discussions. We are grateful to Kerstin Stejskal and Thomas Hummel (Department of Neurobiology, University of Vienna) for providing *Cupiennius salei* and Elvira Lutjanov, Christian Müller, Sabine Ziesemer and Jan-Peter Hildebrandt (all University of Greifswald) for help with western blot analysis. Tim M. Dederichs and Peter Michalik (both University of Greifswald) helped to obtain pictures with the Visionary Digital camera system. Monica M. Sheffer provided helpful linguistic comments that improved the manuscript. This study was financially supported by the German Research Foundation DFG UH 87/11-1 and DFG INST 292/119-1 FUGG, DFG INST 292/120-1 FUGG. POMS was additionally supported by a Bogislaw scholarship from the University of Greifswald and a Laudier Histology cooperation & travel grant.

