## Supplementary material for "Visual pathways in the brain of the jumping spider *Marpissa muscosa*": Fig. S1

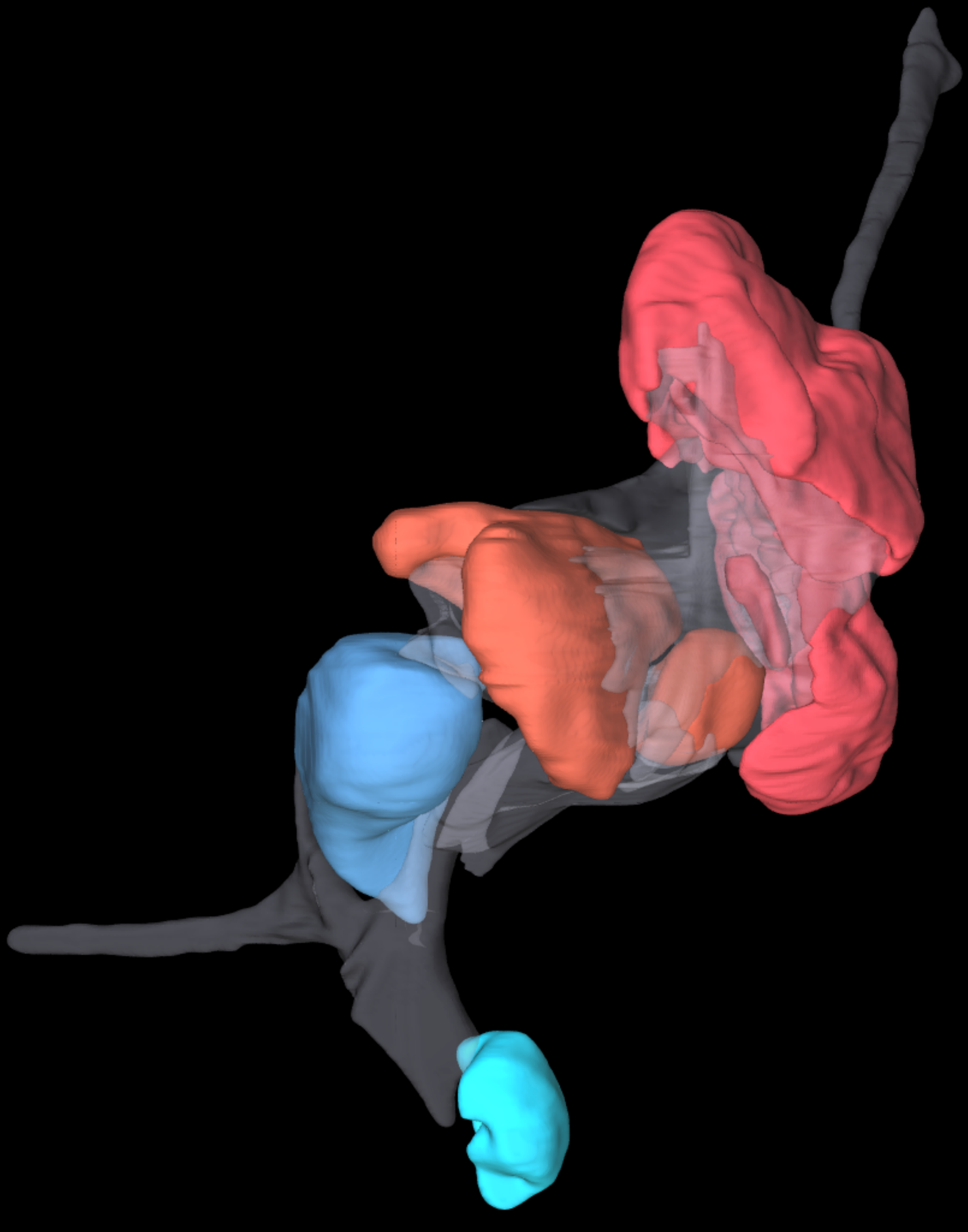

**Figure S1:** Interactive 3D visualization of protocerebral neuropils based on reconstruction of microCT image stacks. The AL1 and the PL1 both are connected to the MG and the L2. A comparatively long optic nerve connects the retina of the PME with its first order visual neuropil, the PM1. **Abbreviations: AL1** first order visual neuropil of the anterior lateral eyes, **L2** shared second order visual neuropil of the anterior lateral and posterior lateral eyes, **MBbr** mushroom body bridge, **MBh** mushroom body haft, **MBp** mushroom body pedunculus, **MG** array of microglomeruli, **PL1** first order visual neuropil of the posterior lateral eyes, **PM1** first order visual neuropil of the posterior median eyes.
